# Decoding Chromatin Dynamics in Cardiac Organoids Reveals Genetic Drivers of Human Heart Development and Disease

**DOI:** 10.1101/2025.10.15.680997

**Authors:** Eyal Metzl-Raz, Ryan Zhao, Salil Deshpande, Yassine Zouaghi, Jackson Powell, Megha Agarwal, Betty B. Liu, Samuel H. Kim, Immanuel Abdi, Ivy Evergreen, Maya U. Sheth, Elizabeth G. Porter, Joshua Rico, Matthew Miyamoto, Julie M. Sanchez, Stephen B. Montgomery, Jesse M. Engreitz, Anshul Kundaje, William J. Greenleaf, Casey A. Gifford

**Author notes:** These authors contributed equally to this work (Eyal Metzl-Raz, Ryan Zhao, Salil Deshpande).

## Abstract

Defining temporal gene regulatory programs driving human organogenesis is essential for understanding congenital defects. We combined a time-resolved, single-cell multi-omic atlas of human induced pluripotent stem cell-derived cardiac organoids with deep learning models of chromatin accessibility, enabling systematic discovery of cis-regulatory syntax underlying heart development and disease. This framework identified cell-state-specific motif syntax, linked motif instances to candidate target genes, and resolved programs governing lineage divergence. Integrating cell-state-resolved molecular profiles with computationally predicted variant effects from congenital heart disease (CHD) cases enabled the prioritization of noncoding variants predicted to disrupt developmental transitions, supporting the paradigm that disease etiology derives from perturbations to regulatory networks governing cardiogenesis. Experimental validation demonstrated that an intronic *ANGPTL2* variant altered differentiation outcomes, implicating *ANGPTL2* in CHD. This study bridges developmental regulation with disease genetics, establishing a framework for discovering the genetic and molecular basis of congenital disorders.

## Introduction

Organogenesis, the formation of functional organ systems, begins in the third week of human development and requires the coordinated differentiation of multiple cell lineages^1–3^. Disruption of the gene regulatory networks that control these cell fate decisions can cause congenital anomalies^4–6^, with congenital heart defects being the most common^7^, affecting up to 3% of live births^8^. Although congenital heart disease (CHD) is heritable, >50% of cases remain genetically unsolved^9–12^. This gap likely reflects the difficulty of resolving the complex genetic architecture of CHD, which has many causal mechanisms that arise during early developmental windows and whose etiology potentially involves multiple coding and non-coding variants acting in tandem^,4,7,13,14^. In particular, cardiogenesis depends on transient progenitor states and coordinated paracrine signaling between nascent lineages^15–18^, processes that are particularly difficult to study directly *in vivo*.

Human induced pluripotent stem cell (hiPSC) differentiation into organoids offers a tractable route by which to model these stepwise, multicellular events^19^. These *in vitro* systems, which contain multiple organ-relevant cell types, allow for the study of the complex cellular interactions and paracrine signaling that shape organogenesis^20–22^. Compared to *in vivo* models, which are challenging to scale and manipulate genetically, and to homotypic cell cultures, which lack multicellular context, organoids provide a tractable and scalable platform for dissecting dynamic gene regulation programs of cell fate and for functional testing of disease mechanisms through targeted genetic perturbations^23,24^.

Predicting and mechanistically interpreting how regulatory genetic variants alter cell-state-specific chromatin states and transcriptional programs remains a significant challenge. Standard assays for mapping chromatin accessibility can nominate candidate regulatory elements and prioritize overlapping non-coding variants^25–28^. However, they lack the resolution at standard sequencing depths to infer the precise combinatorial, spatial arrangements (syntax) of DNA sequence motifs that mediate cooperative binding of context-specific transcription factors (TFs) at these elements^29^. Further, mapping accessibility does not directly quantify how unseen variants affect regulatory activity by disrupting sequence syntax. Deep learning models trained to predict cell-state-resolved chromatin accessibility and gene expression from DNA sequence offer a complementary approach to learn predictive sequence features at base-pair resolution and predict regulatory variant effects^14,29–33^.

Here, we combine a time-resolved single-cell multi-omic (scMultiome) atlas of cardiac organoid differentiation with base-resolution deep learning models to decode cell-state-specific regulatory syntax that drives early human heart development and to decipher non-coding regulatory CHD variants. We profile approximately 120,000 cells across six time points and train neural networks to predict chromatin accessibility from DNA sequence, enabling systematic identification of lineage- and state-specific regulatory rules. This framework nominates motif syntax involving core cardiac regulators, and can distinguish between heterotypic configurations such as HAND+TBX and HAND+TEAD that are associated with divergent fate trajectories. We experimentally validate key predictions by perturbing Myocardin (*MYOCD*), which disrupts ventricular cardiomyocyte specification and reveals non-autonomous effects on ectodermal trajectories, underscoring the importance of paracrine signaling during differentiation. Finally, integrating cell-state-specific chromatin and expression dynamics with model-predicted effects of more than 5 million genetic variants nominates noncoding variants predicted to perturb discrete developmental transitions, supporting a model in which many CHD mechanisms arise during the earliest stages of embryonic development. Ultimately, this work provides a mechanistic framework for decoding early cardiac gene regulation and a resource for interpreting how noncoding variation contributes to CHD.

## Results

### Time-resolved scMultiome profiling of cardioids maps lineage specification and differentiation dynamics

To map the regulatory landscape of early human heart development, we differentiated human induced pluripotent stem cells (hiPSCs) into cardiac organoids (cardioids) by modulating key signaling pathways, including WNT, TGFβ, and retinoic acid^34,35^ (**Fig. 1A**, **S1A**). The resulting cardioids progressed through key developmental milestones, transitioning from pluripotency (day 0; *SOX2*, *OCT4*) through mesendoderm specification (day 1; *TBXT*) to early cardiac mesoderm (day 3.5; *SRF*) and ectodermal states (day 3.5; *NR2F2*). By day 5 and 10, the cardioids had developed into complex, multicellular structures containing cardiomyocytes (*MYL7*), endothelial cells (*PECAM1*), and non-neural ectoderm (*WNT5A*), among other lineages (**Fig. 1A**).

**Figure 1.**
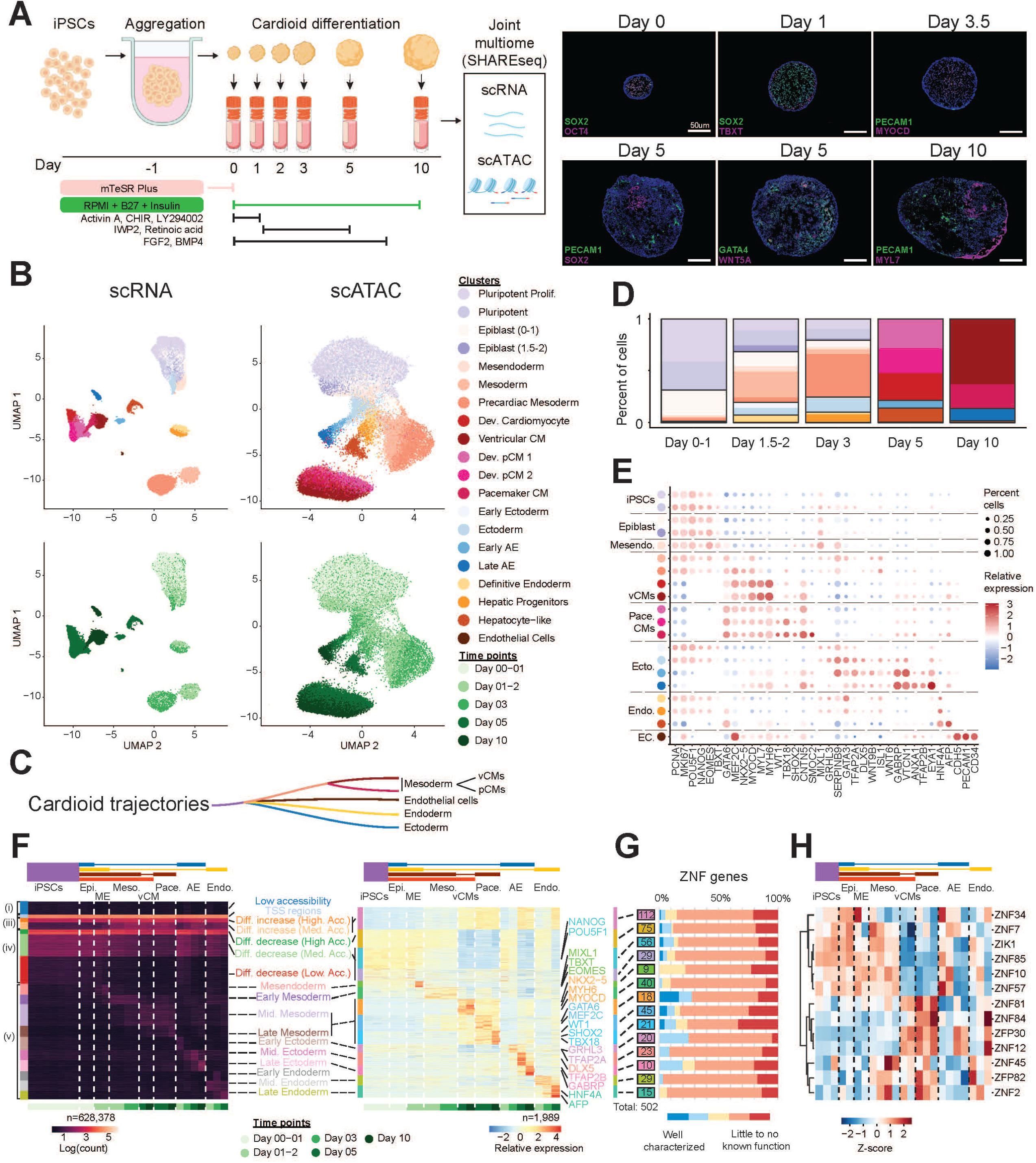
Cardioid scMultiome Atlas Reveals Gene Expression and Chromatin Dynamics. (A) Left: Experimental workflow with the observed lineages in the cardioid. Right: Immunofluorescence staining at multiple differentiation time points. (B) scRNA and scATAC modalities from SHARE-seq multiome processing of ∼60,000 cells throughout cardioid differentiation. (C) Annotated differentiation trajectories identified in the cardioid. (D) Cell type compositions in each sampled day, colors correspond to (B). (E) Manually curated gene markers verify numerous heart-relevant cell types. Colors correspond to (B), z-score normalized per gene. (F) K-means clustering of consensus peaks across all cell types in all time points in the differentiating cardioid (∼630K, left) and k-means clustering of all expressed TFs merged with the top 50 variable genes in each cluster (∼2000 genes). Filled color boxes on top of heatmaps designate trajectories; clusters under the dashed lines are not part of the respective trajectory. (G) Known and predicted function of 502 ZNFs with cell-state specificity as defined in (F) Right. (H) Z-score normalized expression for 13 ZNFs with validated repressor domains.

To resolve the cellular and molecular dynamics of this process, we generated a paired single-cell transcriptome and chromatin accessibility (scRNA/ATAC-seq) dataset from approximately 60,000 cells across six time points (days 0, 1, 2, 3, 5, and 10). We used Seurat^36^ and DecontX^37,38^ for preprocessing, including removal of ambient RNA contamination in the scRNA-seq data, followed by principal component analysis and UMAP embedding (Methods). Clustering analysis identified 20 distinct cell states, revealing a clear temporal progression of lineage specification (**Fig. 1B-E**, **Fig. S1B-D**). Early time points (days 0–2) were dominated by pluripotent (*NANOG*, *POU5F1*), epiblast (*NANOG*, *TBXT*) and mesendoderm (*TBXT*, *EOMES, MIXL1*) populations^34^. These progenitors diversified at intermediate stages (days 3–5) into cardiac mesoderm (*NKX2-5*), non-neural ectoderm (*EYA1*), and hepatic progenitors (*HNF4A*). This transition from proliferative progenitors to differentiated cells (**Fig. 1D**) was further supported by EdU incorporation assays, which showed robust cell division at early stages that progressively decreased over time (**Fig. S1E**). By day 10, the atlas was composed primarily of diverse somatic-like cell types, including ventricular cardiomyocytes (vCMs; *MYL7*, *TTN*), pacemaker-like cardiomyocytes (pCMs; *SHOX2*, *TBX18*), endothelial cells (EC; *PECAM1*, *CDH5*), endoderm-derived hepatic-like cells (Endo; *AFP*, *HNF4A*), and non-neural ectoderm (Ecto; *TFAP2A*, *DLX5*) (**Fig. S1F**). The non-neural ectoderm state exhibited expression of the unique amniotic ectoderm (AE) marker *GABRP*, in addition to *VTCN1* and *TFAP2B*, which supports prior studies suggesting that the epiblast-derived non-neural ectoderm gives rise to AE^39,40^.

Having defined the cellular landscape, we next sought to connect cells’ identities to their underlying regulatory architecture. K-means clustering of 628,378 consensus chromatin accessibility peaks (shared peaks across all cell states out of 9,643,338 total identified peaks in the atlas) revealed dynamic, cell-state-specific patterns. We parsed this data into five main categories of chromatin accessibility: (i) consistently low accessibility in all cell types (n=40,850), (ii) TSS regions (n=11,689), (iii) regions that are either increasing (n=36,270) or (iv) decreasing during differentiation regardless of the lineage (n=173,971), and (v) trajectory-specific accessibility (n=365,598) (**Fig. 1F, left, Fig. S1G**). Accessibility dynamics of these cis-regulatory elements (CREs) broadly tracked the expression dynamics of nearby genes (**Fig. 1F, right, Table S4**). This regulatory association was evident across a core set of 1,989 lineage-defining genes, including transcription factors. The patterns included CREs that were accessible only in specific cell types, as well as those that were dynamic across all lineages during differentiation. Together, these analyses confirm that our cardioid system captures the rapid, lineage-resolved regulatory transitions during early cardiogenesis, providing a high-resolution map connecting transcription factor activity, chromatin state, and gene expression.

### A largely uncharacterized zinc-finger transcription factor program is dynamically regulated during cardioid development

The zinc finger (ZNF) protein superfamily, the most abundant class of transcription factors in the human genome, plays diverse roles as activators, repressors, and architectural chromatin regulators^41,42^. To systematically investigate their potential role in cardiogenesis, we identified 502 ZNF genes whose loci exhibit trajectory-specific chromatin accessibility across 14 developmental stages (**Fig. 1G, Fig. S1H, Table S1**). Using a supervised AI text-mining approach (**Note S5**), we identified a significant knowledge gap: 85% of these dynamically regulated ZNFs are poorly characterized or lack known functions (**Fig. S1H**). Structurally, many of these ZNFs contain domains associated with transcriptional repression, consistent with potential roles in enforcing lineage programs and restricting alternative fates.

Despite the large number of uncharacterized factors, the validity of the mapping of ZNFs to cell states was supported by trajectory-specific activity of well-established cardiac regulators. For example, *ZFPM2* (also called *FOG2*), an essential cardiac progenitor factor and *GATA4* cofactor ^43^ (**Table S1**), was specifically accessible in early and mid-mesoderm clusters, consistent with its established role in very early mesoderm development. Thus, we used the framework to prioritize candidates for further investigation. One example is *CASZ1*, a cardiac transcription factor essential for differentiation and proliferation, with mutations linked to CHD and dilated/arrhythmogenic cardiomyopathy^44–50^, which was specifically assigned to the early developing vCM state (**Fig. S1K**). We also identified *ZNF516*, a mid-mesoderm-expressed KRAB-ZNF reported to suppress EGFR signaling^51^. Together, these examples illustrate how cell-state-resolved chromatin accessibility can connect known regulators to expected developmental windows while surfacing specific, testable hypotheses for under-characterized ZNFs.

To further focus on ZNFs with direct evidence for repression capacity, we examined the subset of 13 ZNFs that contain a functionally validated repressor domain^52^ and exhibited trajectory-specific expression profiles (**Fig. 1H, Fig. S1I-J**). These factors are partitioned into temporally ordered groups, including ZNFs active during early differentiation stages (e.g., *ZNF7*, *ZIK1*), intermediate stages (e.g., *ZNF2*, *ZNF82*), and late stages (e.g., *ZNF84*, *ZFP30*). Additionally, some ZNFs showed lineage-specific activity, such as *ZNF81*, mainly active in pCMs; *ZNF34*, active during later stages of the Endoderm lineage and early differentiation; and *ZNF2*, *ZNF45*, *ZFP82*, which display lower expression in AE. Notably, 12 of these 13 ZNFs do not have any annotated functions. They may represent a potentially novel set of candidate regulators that shape developmental transitions and stabilize lineage identity during human cardioid differentiation.

### Cardioids recapitulate human organogenesis and cardiac lineage specification trajectories

To demonstrate that the cardioid model broadly recapitulates early human development, we projected our scRNA-seq data onto a reference atlas of the human gastrula (**Fig. 2A-B**)^3,36^. This comparison revealed a strong alignment between our *in vitro* cell states and key *in vivo* stages of embryogenesis, including the epiblast, primitive streak, emergent mesoderm, and advanced mesoderm (**Fig. S2A**). Notably, the earliest time points of differentiation (days 0–3) showed the highest mapping scores to the gastrula reference, with the scores decreasing at later stages as cells progressed toward more specified fates (**Fig. 2B**). These results establish that the cardioid system is a suitable model for investigating the early, often inaccessible timeframe of human gastrulation and nascent organogenesis.

**Figure 2.**
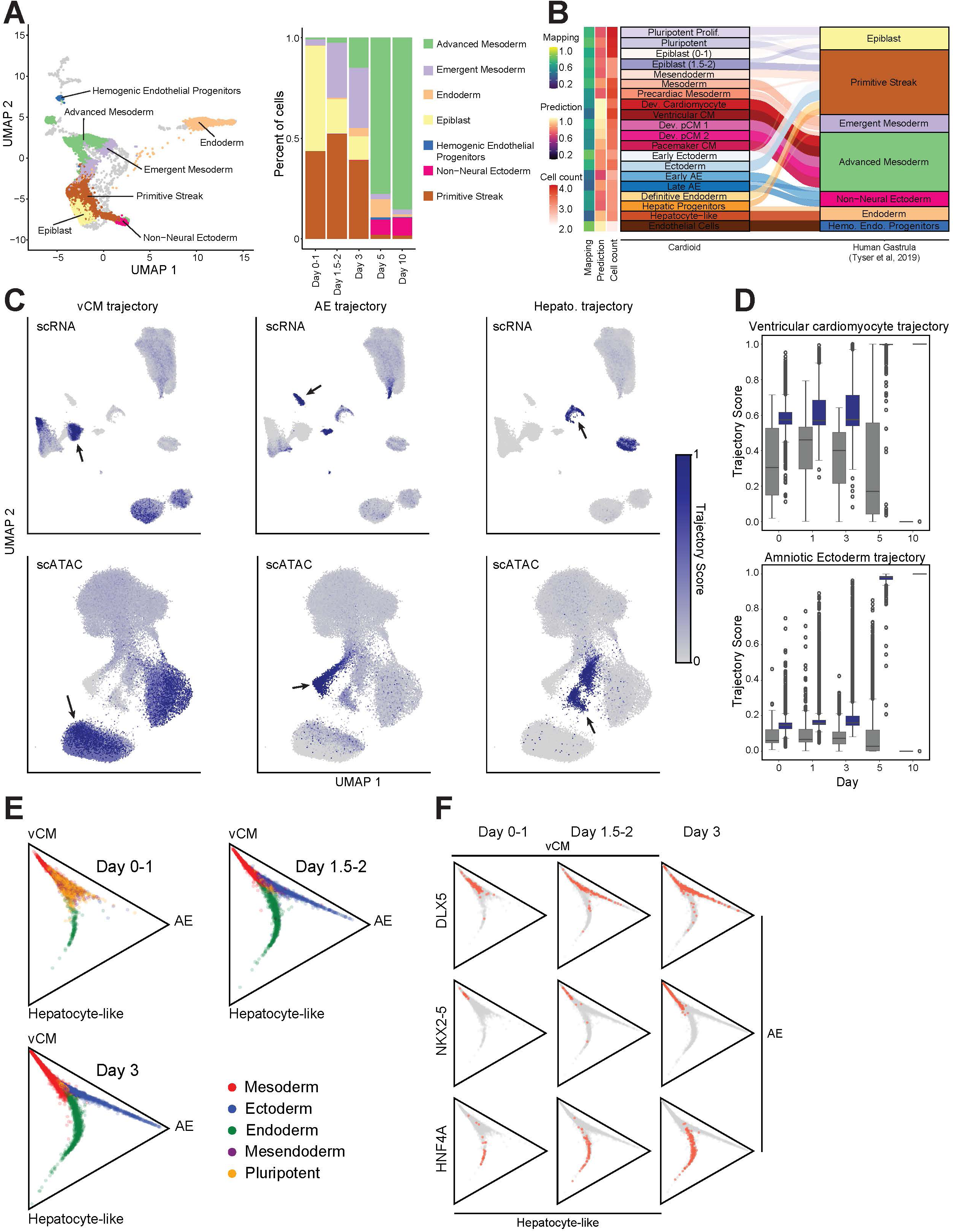
Cardioid Differentiation Occurs Rapidly and Reflects In Vivo Developmental States. (A) Azimuth projection of the cardioid scRNA data to a human gastrula dataset. Left - Mapping the cardioid cells to the human gastrula, UMAP shows overlapping cell states and labels from the human gastrula. Right - earlier cardioid time points’ cells are mapped to earlier human gastrula developmental stages. (B) The projection breakdown shows the mapping of individual cells to the reference. Less mature cell states in the cardioid map have higher Mapping and lower Prediction scores, while more mature cardioid cell states have higher Prediction scores and lower Mapping scores. (C) Optimal Transport Trajectory analysis identifies distinct cells predicted to differentiate to a more mature state (Black arrow). (D) Trajectory scores obtained through OT analysis within predefined trajectories (blue), as shown in Fig.1D, are significantly higher than those outside these trajectories (grey). (E) The Optimal Transport Fate by differentiation day displays the fate probabilities for each cell, highlighting the early commitment of cells toward either of the final states (Late AE, vCMs, Hepatocytes). (F) Optimal Transport Fate. Expression of cell state markers, shown are the top 90% of cells expressing these markers.

Having confirmed the relevance of our model to early embryogenesis, we next sought to reconstruct the developmental trajectories that lead from pluripotent cells to differentiated lineages. To achieve this, we applied Waddington-OT (WOT), a computational approach that infers lineage relationships by calculating the most probable fate of each cell between successive time points^53^. This analysis identified discrete differentiation trajectories (**Fig. 2C, S2B**). For example, we could trace a continuous progression from pluripotency through mesoderm (days 1-2), into pre-cardiac mesoderm (day 3), and ultimately to ventricular cardiomyocytes by day 10.

This lineage inference approach demonstrated that lineage-specific gene expression profiles necessary for proper differentiation are observed within 24 hours of differentiation induction. We observed trajectories associated with each embryonic germ layer, but detected mesodermal trajectories with a higher propensity, which is expected given the differentiation protocol^34^ (**Fig. 2D, Fig. S2C**). By day 2, distinct progenitor populations were already predicted to give rise to ventricular cardiomyocytes (vCMs), AE and hepatocyte-like cells. By day 3, many of these cells showed strong commitment to their terminal fate (**Fig. 2E-F, Fig. S2D**), expressing increasingly mature gene programs (**Fig. S2E**). The trajectory of pCMs shared cell states with the vCM trajectory (Mesoderm and Pre-cardiac Mesoderm), but diverged at day 5 (**Fig. S2B-C**). This suggests a common progenitor state is shared between a pCMs and vCM state as has been observed *in vivo*^54^ and in 2-dimensional hiPSC differentiation^55^. The early lineage specification observed in our model aligns with gene expression signatures observed during human gastrulation *in vivo,* underscoring the model’s capacity to accurately recapitulate gene expression programs that govern cell fate decisions.

### Deep learning models of cell-state-resolved chromatin accessibility decipher dynamic transcription factor motifs regulating cardiac lineage commitment

To decipher the sequence syntax that mediates regulation of chromatin accessibility by TFs defining lineage commitment, we trained ChromBPNet deep learning models to predict base-pair resolution, cell-state-resolved chromatin accessibility profiles directly from DNA sequence. ChromBPNet also corrects distortions in ATAC-seq profiles caused by sequence biases of the Tn5 enzyme used in the experiment^29^. Model interpretation techniques can be applied to assess the predictive contribution of every nucleotide in each regulatory element to the element’s predicted accessibility profile, revealing TF motifs and their syntactic constraints which govern cell-state-specific accessibility patterns^56,57^. This approach overcomes the limitations of traditional motif scanning by learning the quantitative effects of *de novo* discovered motifs on cell-state-resolved accessibility conditioned on their local genomic sequence context.

We first trained cell-state-resolved models for each of the clusters identified across cardioid differentiation (**Fig. 3A**). To evaluate model performance, we computed the Pearson correlation between observed and predicted total counts across peaks for each cell state; correlations ranged from 0.78 to 0.88 (median 0.85) (**Fig. S3A-B**). We also assessed profile accuracy using the Jensen–Shannon divergence (JSD) between observed and predicted base-resolution count distributions within peaks; median JSD values (smaller is better) ranged from 0.47 to 0.73 across models (**Fig. S3A**). As an illustrative example, the ventricular cardiomyocyte model accurately recapitulated the base-resolution accessibility profile at the promoter of *NKX2-5*, a key cardiac regulator (**Fig. 3B, left**).

**Figure 3.**
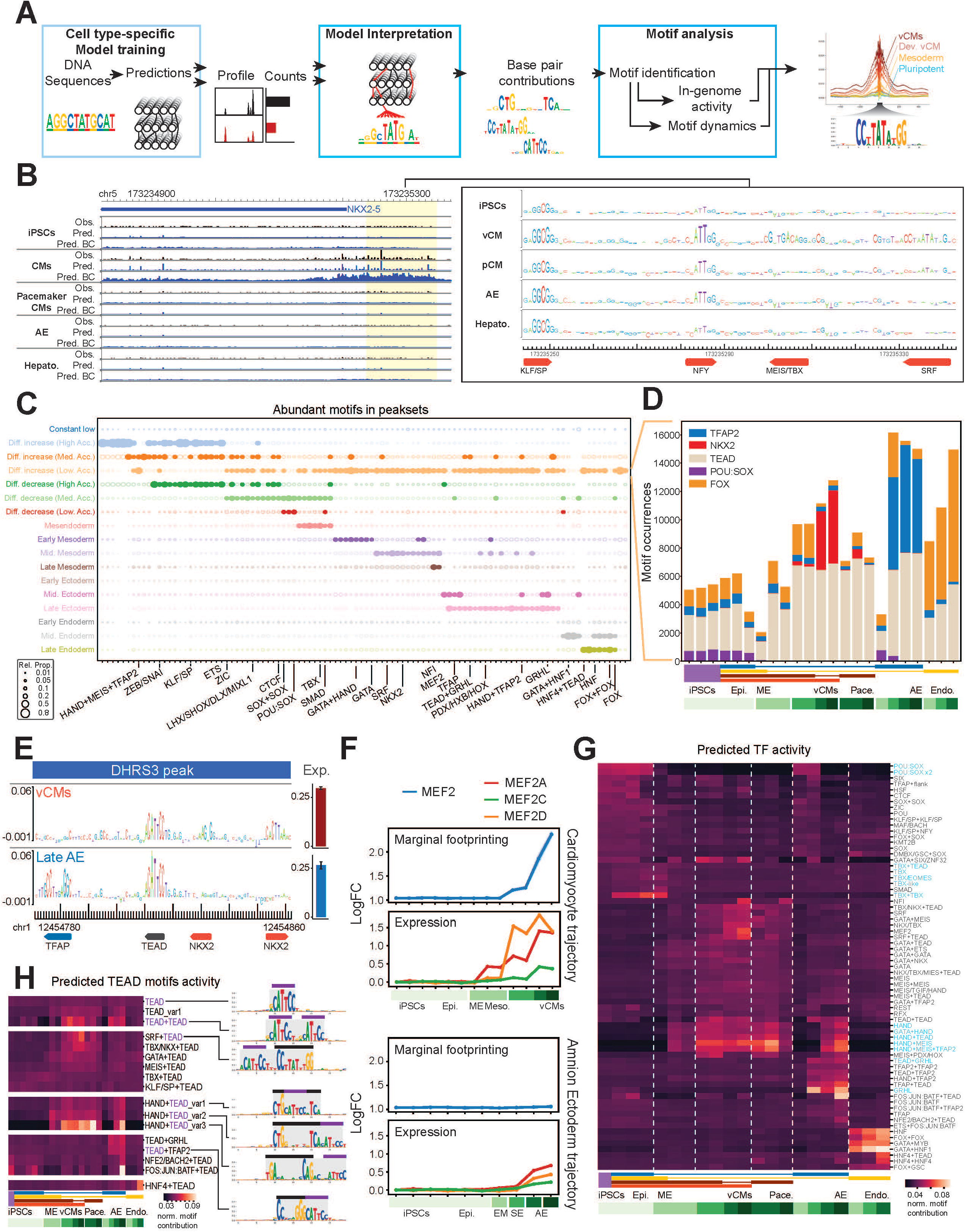
Deep Learning Uncovers Complex Motif Syntax During Cardioid Differentiation. (A) ChromBPNet is a basepair-resolution deep learning model that predicts cell state-specific TF activity. (B) Observed (black) and predicted (before and after bias correction, top and bottom blue tracks, respectively) accessibility in the *NKX2-5* locus for several cell types (left panel) shows increased accessibility and a high correlation with predicted accessibility in mature vCMs. The TSS and proximal promoter regions (highlighted, right panel) are predicted to have high contributions from KLF/SP, NFY, MEIS/TBX and SRF motifs. (C) Performing motif abundance enrichment in peak sets (Fig. 1F) using ChromBPNet predictions allows us to identify specific cell-state profiles of accessible motifs. (D) An allegedly canonical peak set (accessibility increases with differentiation in all lineages) has, in fact, cell-type-specific compositions of different motifs. Whereas TEAD has the same activity dynamics in all lineages in these peaks, other TFs show cell-state-specific activity. (E) Cell-state-specific motifs differentially regulate a *DHRS3*-related peak: Left - TEAD is active in both cell types, but TFAP and NKX2 motifs are cell-state-specific. Right - *DHRS3* log-norm expression. (F) marginal footprinting for MEF2 motif (blue) in two lineages (upper versus lower panels) and corresponding expression dynamics (color lines). In the vCM lineage, MEF2 motif activity is explained by sequential expression patterns of the *MEF2* genes. Although MEF2 is lowly expressed in the AE lineage, it is not predicted to have binding activity. (G) *in silico*, row-normalized cell-type-specific motif contribution (Marginal footprinting) shows various predicted motif activities throughout lineages. Highlighted trends indicated by blue font. (H) The TEAD motif family *in silico*, row-normalized contributions, reveals differential predicted activity of TEAD variant motifs and TEAD composite motifs. Purple bar - TEAD motif; Black bar - partner motif.

We next used the DeepLIFT method to interrogate each model and estimate sequence contribution scores for all peaks in each cell state^56,58^. We then used the TF-MoDISco motif discovery method to infer globally predictive *de novo* sequence features, corresponding to TF motifs, learned by each model^57^. Unifying motifs across models yielded a comprehensive regulatory lexicon of 418 predictive patterns, consisting of 172 distinct motifs from 106 TF families^59^. The majority of motifs were identified as canonical singleton motifs with or without extended predictive flanking sequence (140 and 74, respectively). More than a third of the *de novo* motifs were multi-partite composite motifs, consisting of both heterotypic and homotypic motif combinations (89 and 53, respectively). We also identified 63 motifs of unknown identity, including 43 with positive contributions to accessibility and 20 with negative contributions (**Fig. S3C**). We defined motif families as sets of motifs with highly similar core sequence preferences and variable flanking bases (**Methods**). Cell-state specificity of predictive motifs was evident at the *NKX2-5* promoter, where binding sites for vCM-specific TFs such as MEIS/TBX and SRF showed high contribution scores from vCM models but not from models of other lineages (**Fig. 3B, right**).

We next used the unified motif lexicon to characterize the dynamic activity of globally predictive motifs and their associated TFs across all cell states. Consistent with known lineage programs, motifs for SMAD and TBX were enriched in mesendoderm, MEF2 and NKX motifs in cardiac mesoderm, and HNF and TFAP2 motifs in endoderm and ectoderm, respectively (**Fig. 3C, Fig. S3D**).

Finally, we leveraged cell-state-resolved contribution scores to map predictive motif instances from the unified lexicon across all peaks and cell states and to infer combinatorial TF occupancy patterns (**Methods**). Peaks contained on average eight predictive motif instances, yielding a substantially more specific annotation of active motif syntax than traditional motif scanning, which identified a higher number of motif instances (90 per peak on average) (**Fig. S3E**). Quantifying motif instances across peaks within each cell state revealed both shared and cell-state-specific TF motifs (**Fig. 3C**). Together, these analyses show that ChromBPNet models capture local and global TF occupancy patterns through predictive motif syntax that helps explain accessibility dynamics across the differentiation time course.

### Integrating transcription factor motif and expression dynamics resolves context-dependent regulation of accessibility by distinct TF family members

The precise regulation of gene expression is governed by a complex interplay between TFs and cis-regulatory elements (CREs). A gene’s expression can be regulated by different TFs binding to multiple distinct CREs, and/or by different TFs acting on the same CRE in a cell-state-specific manner. The latter, which we refer to as context-dependent cis-regulatory elements (cdCREs), are particularly challenging to decipher. Our integrative framework can resolve both types of regulatory logic. The cdCRE upstream of *DHRS3*, a gene involved in regulating retinoic acid levels during embryogenesis^60^, serves as a clear example. While this single element is accessible in multiple lineages, our models predict that distinct TF occupancy patterns contribute to its accessibility in different cell contexts, via lineage-specific motif contributions within the same CRE (**Fig. 3E**). In vCMs, NKX2 motifs are predicted to regulate accessibility. In contrast, in AE, *TFAP2* motifs are predicted to regulate accessibility, while TEAD motifs contribute similarly in both cell states. Examining cell-state-resolved expression of TFs from the NKX family nominates the most likely TFs binding these predicted motifs. In the vCMs, *NKX2-5* is the highest expressed NKX family member, aligning with its known role as a critical regulator for proper heart development. In AE, other members of the NKX family are expressed (*NKX2-3*, *NKX2-6*, *NKX3-1*), although the models do not predict NKX family motif contributions to *DHRS3* accessibility in AE (**Table S12**).

We extended this analysis to examine a broader set of peaks accessible across multiple terminal lineage states (**Fig. 3C**, light orange fourth row). Despite exhibiting similar accessibility across these cell states, the most prevalent predicted drivers of these peaks were lineage-specific. For example, the NKX2 motif family was the most prevalent motif in these peaks in vCMs and was not a key driver in any other lineage. Similarly, TFAP2 was the largest driver in and specific to AE, OCT:SOX was specific to hiPSCs and early states across all lineages, and FOX showed the highest presence in endoderm-derived cells (**Fig. 3D, Fig. S3F**). These motif dynamics are consistent with lineage-specific TF expression patterns (**Table S13**). These results highlight the models’ ability to disentangle the regulatory logic underlying accessibility, suggesting that cdCREs with similar accessibility across cell states may encode pleiotropic motif syntax reflecting context-dependent regulation by distinct TFs.

Our framework can also resolve the ambiguity of multiple TFs from the same family binding to similar motifs. The FOX family, which contains many members implicated in CHD, illustrates this challenge. While the canonical FOX motif is active in nearly all cell states, by integrating our models’ FOX motif predictions (**Fig. 3D**) with single-cell gene expression (**Table S13**), we can nominate the specific FOX family members responsible for this activity in a cell-state-specific manner. For example, we find that the CHD-associated TFs *FOXP1* and *FOXF1* are specifically expressed in vCMs, whereas other family members like *FOXM1* and *FOXO1* are transcriptionally inactive in the same cells (**Table S13**). These examples demonstrate our framework’s ability to link transcription factor expression to motif activity in a cell-state-specific manner.

When comparing cell-state-specific motif proportional composition, different cell states might show similar composition of attributing motifs (**Fig. S3G**, top). However, even cell states with similar motif compositions exhibit large differences in the absolute number of motif instances (**Fig. S3G**, bottom). This suggests that the proportional usage of the regulatory motif lexicon is partly conserved across developmental lineages. Together, these results support a model in which cell states draw on a shared motif lexicon, but deploy it in distinct combinations and proportions to achieve lineage-specific regulatory programs.

### *In silico* marginalization disentangles the effects of individual motifs and cooperative motif syntax on context-dependent accessibility

A central goal in developmental genomics is to resolve how TFs contribute to gene regulation within complex, multi-motif enhancers and across cell states. To address this, we used an *in silico* marginalization strategy^61,62^. This technique estimates a motif’s impact on chromatin accessibility within each cell state by comparing model predictions on background genomic sequences with and without the motif inserted, thereby isolating its marginal effect independent of the confounding influence of neighboring motifs in accessible regions. Furthermore, this allows us to quantify the marginal influence of rare motifs on cell-state-specific accessibility, which are often overlooked by conventional methods that rely on statistical enrichment.

The *MEF2* family provides a clear case study. While MEF2 motifs are abundant in mesodermal peaks and several MEF family members (*MEF2A*, *MEF2C*, *MEF2D*) are expressed in both vCM and AE lineages, our analysis revealed that *MEF2*-driven accessibility is largely restricted to the vCM lineage. We predict a wave of MEF2 accessibility that begins in pre-cardiac mesoderm and peaks in vCMs, which correlates with the sequential expression of *MEF2A*, *MEF2D*, and *MEF2C* (**Fig. 3F**, upper panel). In contrast, our model does not predict MEF2 motif accessibility in the AE lineage despite detectable expression of *MEF* genes (**Fig. 3F**, lower panel). This demonstrates our approach’s ability to discriminate between TFs that are expressed and those that are actively shaping the chromatin landscape in different cell contexts.

By extending *in silico* marginalization to pairs and higher-order combinations of motifs, we can decipher the effects of cooperative TF binding on chromatin accessibility. The model predicts, for instance, that a TBX homotypic motif has strong synergistic effects on accessibility compared to a TBX monomer during organogenesis-related stages, while a tandem OCT:SOX motif has no additional effect on accessibility over a monomer in pluripotent stem cells (**Fig. 3G**, **Fig. S3H**). Conversely, a *GRHL* monomer has stronger predicted activity compared to GRHL+TEAD heterotypic syntax (**Fig. 3G**). We also observed a complex HAND code whereby tandem, heterotypic syntax with multiple TFs (e.g. GATA, TEAD, MEIS) exhibits predicted regulatory activity in both cardiomyocytes and the AE, except for HAND+TFAP2 instances which are predicted to be exclusively active in the AE (**Fig. 3G**).

The TEAD motif provides an instructive case study in predicted context-dependent regulatory syntax. While the core TEAD motif is present in accessible CREs across most cell states, its contribution to accessibility becomes highly cell-state-specific in the proximity of motifs of other TFs (**Fig. 3H, Note 4**). Specifically, TEAD is predicted to cooperate with SRF motifs in cardiomyocytes, with GRHL in the ectoderm, and with HNF4 in the endoderm. The model’s precision extends even to the precise spacing constraints between motifs. For example, the HAND and TEAD motif pair has the highest effect of accessibility when spaced exactly three base pairs apart, demonstrating the model’s ability to precisely resolve subtle syntactic constraints that define lineage-specific enhancer accessibility (**Fig. 3H**). Other prominent TF families exhibit the same sequence-specific activity patterns (**Note 4**). Together, these results show that the models not only predict which motifs are active, but also how their combinatorial arrangements and precise spacing syntax encode cell-state-specific regulatory logic.

### Integrating peak-gene links with accessibility-derived motif syntax prioritizes TF regulators of developmental gene programs

To link accessibility-derived motif syntax into gene-level regulation, we first used the scE2G method^63^ to infer associations between 628,378 consensus peaks to putative target genes, which resulted in 1,256,939 quality-controlled peak-gene links across all cell states (**Fig. S4A**, **Table S15**). To estimate each motif’s predicted influence on a gene in a specific cell state, we then summed contribution scores of all predictive instances of the motif identified by the cell-state-specific accessibility model over all peaks in that cell state linked to the gene (Methods) (**Fig. 4A**, **Fig. S4B**).

**Figure 4.**
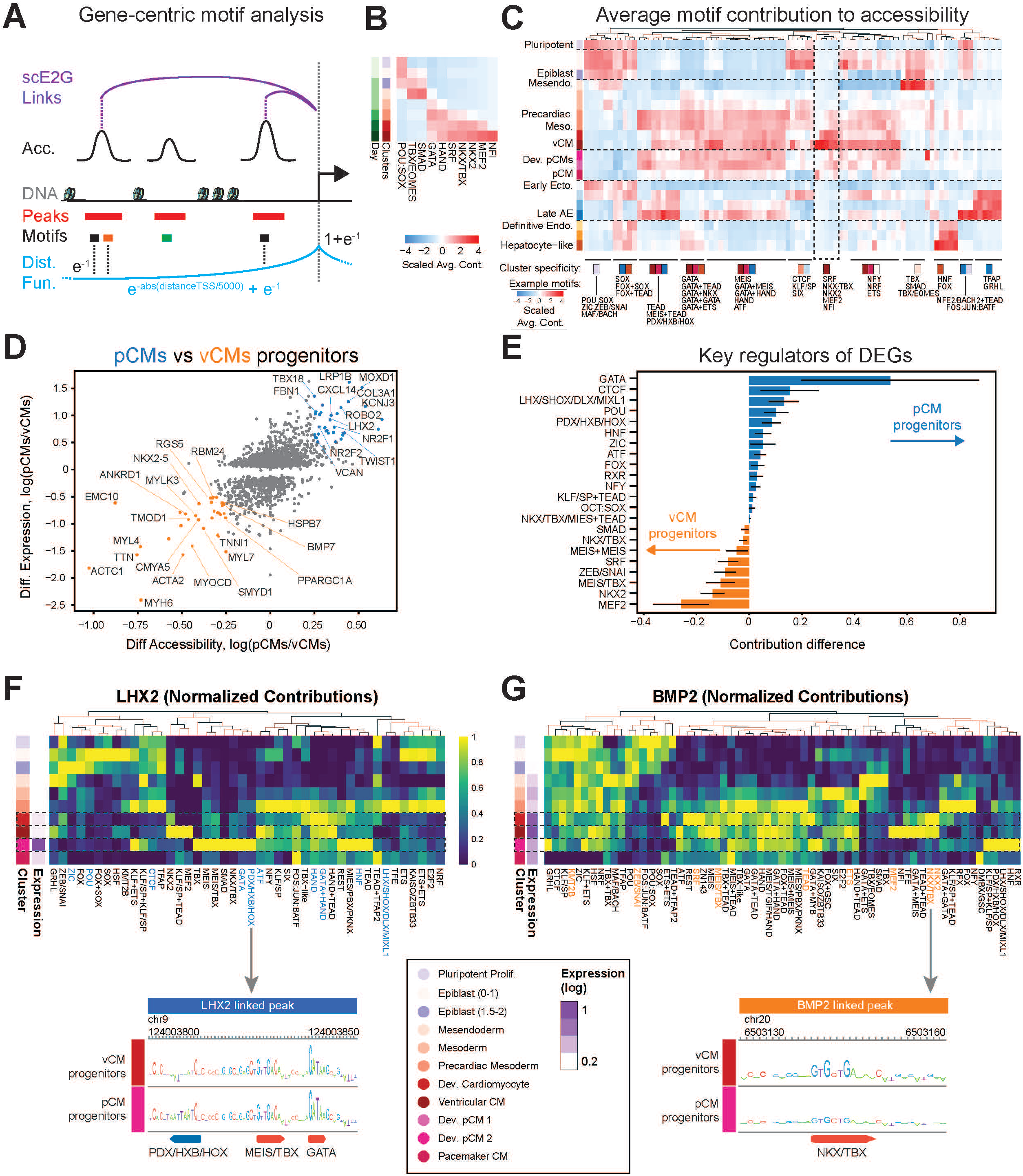
Linking Accessibility to Target Genes Defines Lineage-Specific Gene Regulatory Networks. (A) The gene-centric approach links active motifs in a cell-state-specific manner. Motifs in called peaks are filtered by consensus scE2G peak-gene linkages and are weighted down by a distance function before summing their contributions. (B) Average in genome contribution for the SRF regulome in the vCM lineage. (C) Average in genome contribution for the entire motif compendium across all cell states. The sums of contributions (intensity) for each motif family were averaged and scaled across all expressed genes (approximately 12,000 genes) in each cell state. The major cell-state specificity is indicated at the bottom, alongside motif examples (Fig. S4B for all annotations). (D) Comparison of differential genes (y-axis) versus differential accessibility (x-axis) between vCMs and pCMs progenitors. Marked genes represent the top differentials in either trajectory. (E) Differences in average contributions for each motif between vCM and pCM progenitors. Positive values indicate higher contributions in pCMs, while negative values indicate higher contributions in vCM progenitors. Error bars represent standard deviation (SD). (F-G) Motif syntaxes for *LHX2* (F) and *BMP2* (G), which are differentially expressed genes in pCMs and vCMs, respectively. Dashed boxes highlight vCM progenitors (top) and pCM progenitors (bottom), and the blue/orange annotations indicate the key regulators of the lineages shown in (E). Panels at the bottom display example peaks and motif contributions for each gene.

We first applied this gene-centric motif ranking approach to reconstruct the well-characterized regulatory cascade of vCM specification (**Fig. 4B**). To estimate overall regulatory contributions of motifs for this differentiation trajectory, for each cell state, we averaged the contribution of each motif over all lineage-defining genes (**Fig. 1E, Table S11**). The resulting dynamics of these gene-set level motif contributions across cell states accurately recapitulate known sequential regulatory changes over developmental time: from pluripotency factors (POU, SOX^64^) in hiPSCs, specification-associated TFs (TBX, EOMES, SMAD^65^) in epiblast-like cells, to mesendoderm-associated TFs (GATA, HAND, NKX2^66^) in early progenitors, and finally to the canonical cardiomyocyte TFs (SRF and MEF2^67^) that orchestrate later stages of differentiation *in vivo*. The fidelity of this reconstruction to established principles of vertebrate heart development supports the validity of our gene-centric motif ranking framework.

Expanding this analysis genome-wide and across differentiation trajectories, we generated a global map of gene-centric motif contributions across all cell states (**Fig. 4C**, **Fig. S4B, Table S11**). This map captures the relative contributions to accessibility of both transcriptional activators and repressors, such as *ZEB1/2*^68^ and *SNAIL2*^69^. These repressors are uniquely expressed in the mesoderm lineage, where our models assigned their motifs negative contribution scores, consistent with their role in compacting chromatin^70^. Concordantly, canonical epithelial markers that are silenced during the epithelial-to-mesenchymal transition (EMT), including *CDH1*, *EPCAM*, *CRB3*, and *CLDN*^71^, were all downregulated in the mesoderm, consistent with reduced ZEB/SNAI associated accessibility at their CREs. Additionally, genes known to be activated by ZEB/SNAI, such as *ACTA2*, *VIM* and *FN1*, were upregulated (**Table S14**).

### Inferring regulatory programs distinguishing ventricular and pacemaker cardiomyocytes

Resolving the regulatory programs that differentiate recently diverged cell states has been a major challenge. Our framework addresses this by systematically linking quantitative effects of TF motif syntax driving accessible CREs to the putative target genes in each cell state. By applying this framework to dissect the TF motif syntax distinguishing ventricular (vCM) and pacemaker (pCM) cardiomyocytes, we can directly investigate the subtle differential regulation of critical disease-associated genes in these cell states. To identify the TFs driving this divergence, we selected genes that were both differentially expressed between vCM and pCM progenitors and linked to differentially accessible peaks (**Fig. 4D**), then calculated the difference in each TF motif’s cumulative contribution score across this gene set between the two progenitor states (**Fig. 4E**).

Our analysis revealed that pCM progenitors exhibit high predicted contribution from the GATA, PDX/HOX, and LHX/SHOX motif families. These motifs were linked to key pacemaker identity genes, including *FBN1* and *TWIST1*, which are associated with congenital syndromes^72–75^, and also predicted to regulate highly accessible enhancers of additional pCM-specific genes, including *LHX2*, *TBX18* and the ion channel *KCNJ3* (**Fig. 4F, Fig. S4D, Fig. S4E**). In contrast, vCM progenitors exhibited strong contributions from MEF2, NKX2-5, and SRF motifs, linked to vCM-specific targets like *BMP2* (**Fig. 4G, Fig. S4F**), as well as *TTN*, *ANKRD1* and *MYH6*, which are linked to various cardiomyopathies (**Table S11**). Together, these findings establish that vCM and pCM progenitors are governed by distinct, clinically relevant regulatory syntax, and demonstrate the ability of our framework to resolve the regulatory logic underlying closely related but divergent cell fate decisions.

### CRISPR interference validates predicted ventricular cardiomyocyte regulatory syntax

Based on our model’s prediction that the SRF network is a key vCM-specific driver (**Fig. 4B-C**), we sought to functionally understand the dysregulation mechanism upon genetic perturbation. Given that SRF is a ubiquitously expressed DNA binding cofactor that facilitates MYOCD-directed differentiation events during cardiomyocyte specification^67^, we used a CRISPR-interference (CRISPRi) system to knock down (KD) *MYOCD* to test whether the model’s predictions were specific to MYOCD-SRF interactions (**Fig. 5A-C; Fig. S5F**). *MYOCD* KD resulted in a significant reduction in vCMs by day 10 without affecting overall organoid size or total cell number (**Fig. 5B**, **Fig. S5A**), suggesting a cell fate switch rather than a defect in viability or proliferation. To identify the molecular dynamics affected by *MYOCD* KD, we generated scMultiome data from control-matched knockdown cardioids at days 0-1, 1.5-2, 3, 5, and 10. We observed distinct alternative cell states, specifically in the vCM and AE lineages, where *MYOCD* KD resulted in alternative developing CM cells (**Fig. 5D**). In the AE lineage, we observed a slight shift in the Early AE state, resulting in the lack of a late AE subpopulation (**Fig. 5D**). To test if *MYOCD* KD led to an alternative developmental trajectory, we projected control and knockdown cells onto a murine embryonic development atlas encompassing E8 to P0 embryos^76^. This revealed a qualitative divergence beginning on day 3 (**Fig. 5E; Fig. S5D-E**). While control mesodermal cells progressed toward a first heart field (FHF) ventricular and atrial cardiomyocyte state, *MYOCD* KD cells represented a progenitor state more similar to the second heart field (SHF) (**Fig. 5E; Fig. S5B**). Additionally, projecting the control and *MYOCD* KD cells onto the mouse developmental atlas’s fetal time revealed that *MYOCD* KD CMs exclusively mapped to the E8-E8.5 stage, whereas control CMs mapped mainly to the later developmental stages of E18.5-P0 (**Fig. S5C**). This confirms that *MYOCD* is essential for proper FHF-derived cardiomyocyte specification and that its loss causes a developmental arrest, extending previous *in vivo* observations^67^.

**Figure 5.**
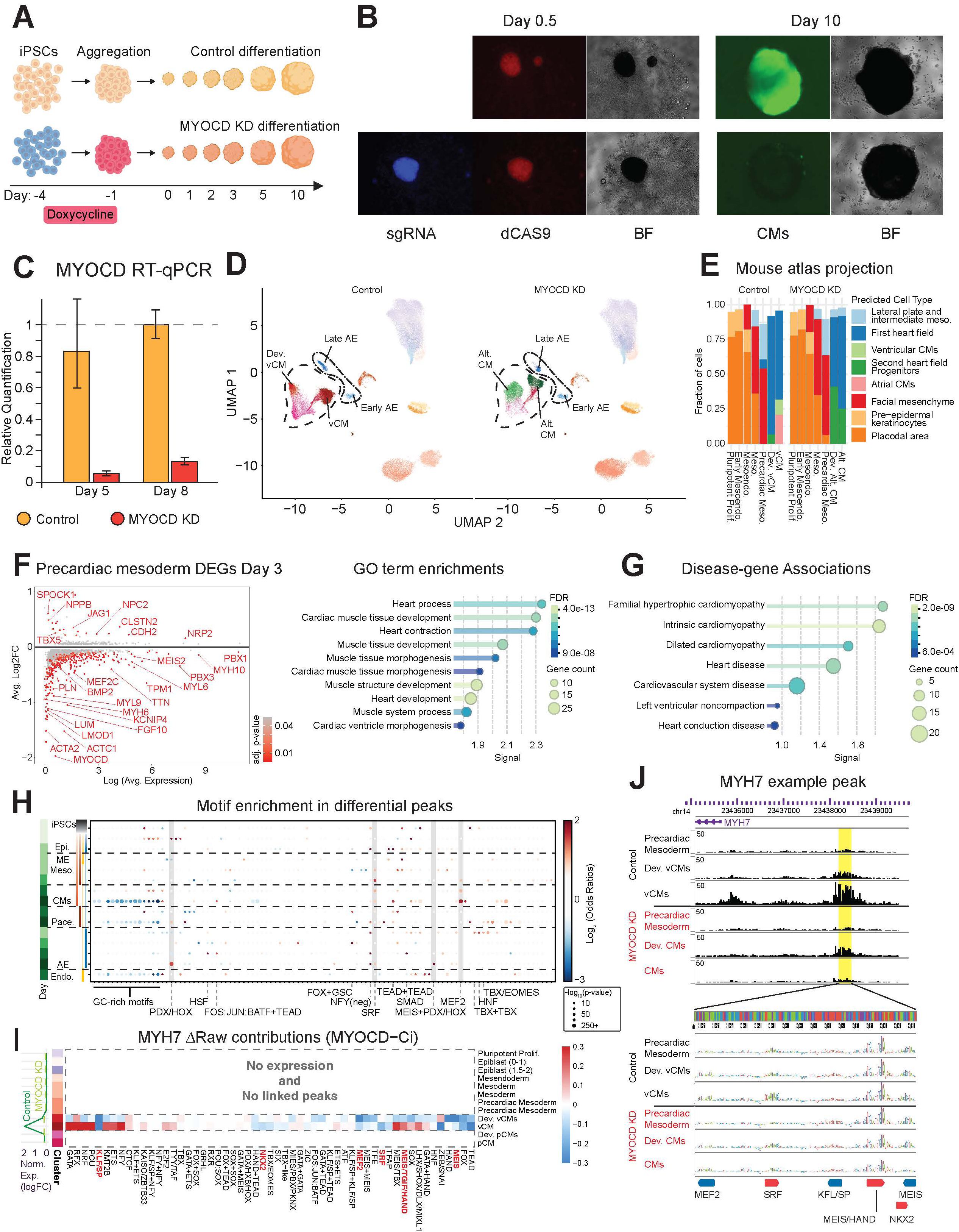
CRISPR Interference Validates Network Predictions (A) *MYOCD* KD experimental scheme. (B) *MYOCD* KD images taken 12 hours after aggregation and on Day 10 of differentiation. On Day 0.5, both the sgRNA (BFP) and the dCAS9 (mCherry) are still visible. By Day 10 of differentiation, the control cardioid displays robust beating vCMs (*MYL7*-*GFP*), while the *MYOCD* KD predominantly lacks them. (C) RT-qPCR relative expression of *MYOCD* in control and *MYOCD* KD cardioids on days 5 and 8 indicates ∼90% KD efficiency. (D) scRNA UMAPs show the depletion of specific cell types (vCMs and AE lineages), whereas other clusters are shared across conditions. (E) Differential prediction of cell types, achieved by projecting the cardioid scRNA data onto a mouse developmental atlas, reveals that alternative vCMs are more closely aligned with an SHF state than with the mature states of CMs. (F) Differential gene expression at Day 3 of cardioid differentiation between *MYOCD* KD and control (left panel) and the associated GO term enrichments (right panel). (G) Disease association of the differential genes determined by DISEASES enrichment analysis^126^. (H) Motif enrichment in differential peaks following *MYOCD* KD. Motifs enriched in peaks with decreased accessibility have positive log odds ratios, whereas those in peaks with increased accessibility have negative log odds ratios following *MYOCD* KD. Grey vertical bars highlight motif examples with high cell-state-specificity. (I) The difference in the raw sum of contribution scores per motif in all *MYH7*-linked peaks in *MYOCD* KD versus control. The dashed box indicates no significant expression and no linked peaks via scE2G in specific cell states. Expression (Dark/light green for Control/*MYOCD* KD, respectively) was normalized to the iPSCs’ basal expression levels. Red motif annotations correspond to motifs identified in the peak example (J). (J) The model contribution scores in a *MYH7*-linked peak upstream of the gene illustrates that decreased accessibility of the peak is attributable to the reduced impact of specific TFs on chromatin accessibility.

Differential gene expression analysis (**Table S10**) pinpointed the timing of the molecular effects induced by *MYOCD* KD. Prior to day 3, the only significantly altered gene was *MYOCD* itself (**Fig. S5G**, “mesoderm”). However, by day 3, significant transcriptomic changes appeared almost exclusively in the precardiac mesoderm (**Fig. 5F, left; Fig. S5G)**. These changes included a significant downregulation of genes associated with heart development (**Fig. 5F, right**), directly linking the perturbation to the molecular programs implicated in heart disease (**Fig. 5G)**.

To identify the core regulatory networks disrupted by *MYOCD* KD, we analyzed motif enrichment within chromatin regions that lost or gained accessibility upon KD (**Fig. 5H**). For each cell state, we performed *k*-means clustering (*k*=3) based on the log_2_ fold change (log_2_FC) between Control and *MYOCD* KD in that specific cell type, aiming for three major chromatin accessibility profiles: (i) decreased, (ii) unchanged, or (iii) increased accessibility upon knockdown. Then, for each motif family, we used Fisher’s exact test to assess motif enrichment between the clusters that gained and lost accessibility post KD, considering only active peaks (containing at least one motif within +/- 500bp of the peak summit). This analysis revealed an apparent dichotomy in the cardiomyocyte lineage - regions that gained accessibility were enriched for general promoter GC-rich motifs implicated in cell states that precede cardiomyocytes, likely reflecting indirect gene regulation of the observed arrested developmental state (**Fig. 5H**). In contrast, chromatin regions that lost accessibility were most significantly enriched for SRF and MEF2 motifs (**Fig. 5H**, grey vertical bars). This result provides direct evidence that the primary autonomous effect of the perturbation is the targeted disruption of the MYOCD-SRF/MEF2 regulatory network, which is essential for cardiac development^67^.

To discern how *MYOCD* KD leads to repression of its target genes, we examined the regulatory syntax at the cardiomyocyte-specific gene, *MYH7*. Our model revealed a significant disruption of the regulatory program at the onset of *MYH7’s* expression. In control cells, *MYH7* is driven by the coordinated activity of various TFs(e.g., MEIS, MEF2, HAND and SRF). However, upon *MYOCD* KD, key activators like MEIS and MEF2 lose their predicted activity at *MYH7* regulatory elements despite remaining expressed (**Fig. 5J**). This regulatory imbalance provides a mechanistic view of how a gene’s regulatory landscape is altered upon MYOCD KD.

Intriguingly, this analysis also suggested *MYOCD* is necessary for the AE lineage, which exhibited significant transcriptomic changes upon KD (**Fig. 5D, Table S10**). Our analysis revealed that the canonical *MYOCD*-associated motifs SRF and MEF2 did not directly drive the disruption in this lineage, but instead, the most significant perturbation was in the activity of the PDX/HOX transcription factor family (**Fig. 5H**). The indirect effect of *MYOCD* KD on PDX/HOX motifs (**Fig. S5H**) suggests that defects in cardiomyocyte lineage differentiation may lead to cell non-autonomous perturbed signaling in the AE lineage in the cardioid. Indeed, our data indicate that *MYOCD* KD reduces *BMP2* expression in cardiomyocytes, a morphogen critical for AE differentiation^77^ (**Fig. S5I**). Given that BMP signaling is a known inducer of *HOX* gene expression^78–80^, these findings suggest a paracrine mechanism whereby aberrant BMP signaling derived from the perturbed mesoderm prevents faithful AE specification in vitro, where it converges on the PDX/HOX regulatory network to mediate the non-autonomous effect. Collectively, these data demonstrate the capacity of deep learning frameworks integrated with organoid systems to resolve the precise regulatory syntax underlying multicellular developmental transitions that are intractable to direct *in vivo* investigation.

### Cell-state-specific gene regulatory programs of genes associated with congenital heart disease

To identify cell states disrupted in CHD, we evaluated the gene expression pattern of 247 genes implicated in CHD^81^. We first combined them into expression modules via k-means clustering (**Fig. 6A, Fig. S6A-B**), which revealed that 97% of genes are expressed across multiple lineages in our data. This broad multi-lineage expression highlights the need to consider diverse early cell states alongside somatic cell types when interpreting CHD variants and designing functional validation experiments.

**Figure 6.**
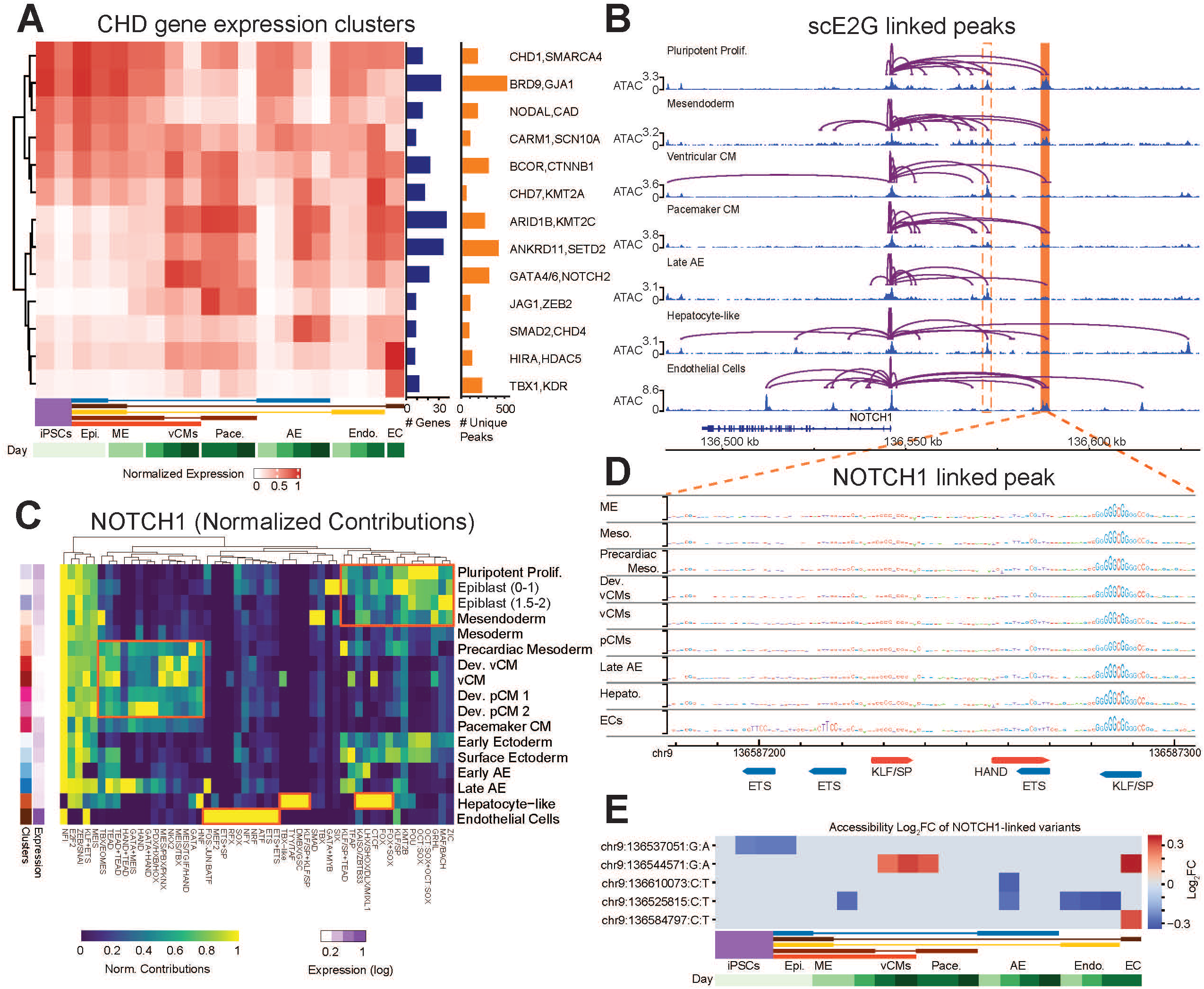
CHD-associated Gene Regulatory Networks Captured in Cardioids. (A) 247 CHD genes clustered (via *K*-Means, n=17) by normalized gene expression across cardioid cell states. Blue bars are the number of genes in the cluster. Orange bars are the number of unique linked peaks for all genes in the cluster. (B) *NOTCH1*-linked peaks in selected cell states. For each cell state, shown are linked peaks in purple and the corresponding ATAC tracks. The orange bar highlights a cdCRE element zoomed in (D). The dashed orange lines indicate an additional peak highlighted in Fig. S6F. (C) *NOTCH1* motif contributions. Orange boxes indicated trends highlighted in text. Column normalized. (D) Motif contributions in a *NOTCH1*-linked cdCRE peak (highlighted in (B)). (E) Five gnomAD rare variants found to disrupt accessibility in *NOTCH1*-linked peaks.

To resolve regulatory logic for the genes implicated in CHD^81^, we next integrated cell-state-specific motif derived from ChromBPNet models with scE2G links. This analysis identified 2,262 unique putative CREs linked to known CHD genes, with a median of 5 linked CREs per gene (**Fig. S6C**). *NOTCH1* is a prominent CHD gene associated with myriad structural defects of disparate origins^82–85^, and was expressed early in differentiation in all lineages as well as the EC state (**Fig. S6D**). Beyond its well-studied role in endothelial cells^86^, *NOTCH1* is also required for second heart field development^87^, neural crest-derived smooth muscle differentiation^88^ and the diversification of cells derived from the pharyngeal pouch endoderm^89^ into FGF-secreting cells critical for outflow tract and aortic arch development in the heart^90^. This multi-lineage role of *NOTCH1* makes it an informative test case for our framework. Indeed, we identified 28 unique peaks predicted to regulate *NOTCH1* across all cell states, highlighting cell state-specific putative CREs (**Fig. 6B**).

Given our aforementioned observations that peak level analyses obscure TF-driven regulatory logic, we next defined the cell-state-specific *NOTCH1*-associated motif syntax based on ChromBPNet nominations with scE2G-linked peaks (**Fig. 6C, Fig. S6E**). This analysis defined the TF families that contribute to accessibility along the differentiation trajectories. These motif families correlated with states displaying graded *NOTCH1* repression, demonstrating the capacity of distinct motif family architectures to govern *NOTCH1* expression. Various patterns were predicted to collectively regulate *NOTCH1* in different cell states. For instance, during early differentiation stages, OCT:SOX motifs constitute the principal contributors, whereas in endothelial cells, where maximal *NOTCH1* expression was observed, ETS, SOX, NFY, and MEF2 motifs, in addition to the ETS-ETS tandem homotypic configuration, demonstrate the greatest contribution (**Fig. 6C-D**).

To identify regions that encode human variants that may disrupt *NOTCH1* regulation, we next quantitatively evaluate the effect of 97 rare (MAF < 0.001) single-nucleotide variants (SNV) from gnomAD^91^ (**Table S16-S17**) encoded within regions predicted to interact with the *NOTCH1* promoter. These analyses revealed two variants predicted to exert cell-state-specific effects on accessibility (**Fig. 6E**), supporting the motif nominations. This result highlights a key distinction between coding and noncoding variation. Whereas coding variants typically affect all isoforms and expressing cell types, noncoding variants in CREs can disrupt gene expression selectively in specific cell states, potentially producing phenotypes that differ from or are altogether absent in individuals carrying coding variants in the same gene.

### Noncoding variants in congenital heart disease exhibit cell-state-specific regulatory effects in cardioid trajectories

CHD encompasses a broad spectrum of structural malformations present at birth. While the majority of genetic etiologies remain incompletely defined^7^, established mechanisms involve perturbations of the nascent stages of cardiogenesis - an embryonic phase that is challenging to characterize and experimentally validate due to the limited availability of physiologically relevant model systems^9,92,93^. This challenge is especially pronounced for noncoding variants, which are predicted to exert effects in a highly cell state–specific manner^14,30,31,92–95^.

To identify *de novo* and inherited noncoding variants that may contribute to CHD etiologies, we next used our cell-state-specific ChromBPNet models to predict the effects of single nucleotide variants (SNVs) and short indels from CHD probands (-derived)^9^ and gnomAD^91^ on local chromatin accessibility. In total, we predicted the effects of over five million variants across 29 cardioid cell states (**Fig. 7A, Table S16-S17**). Across allele frequencies, 2-5% of evaluated variants were prioritized by at least one cardioid ChromBPNet model (**Methods**). Approximately 60% of prioritized variants impact a defined DNA sequence motif within our atlas. Notably, 16% of CHD-derived *de novo* variants that impact motifs were encoded within scE2G-linked peaks, compared to only 5-7% of gnomAD variants, suggesting enrichment of *de novo* CHD variants in CREs with predicted gene regulatory activity (**Fig. 7A**).

**Figure 7.**
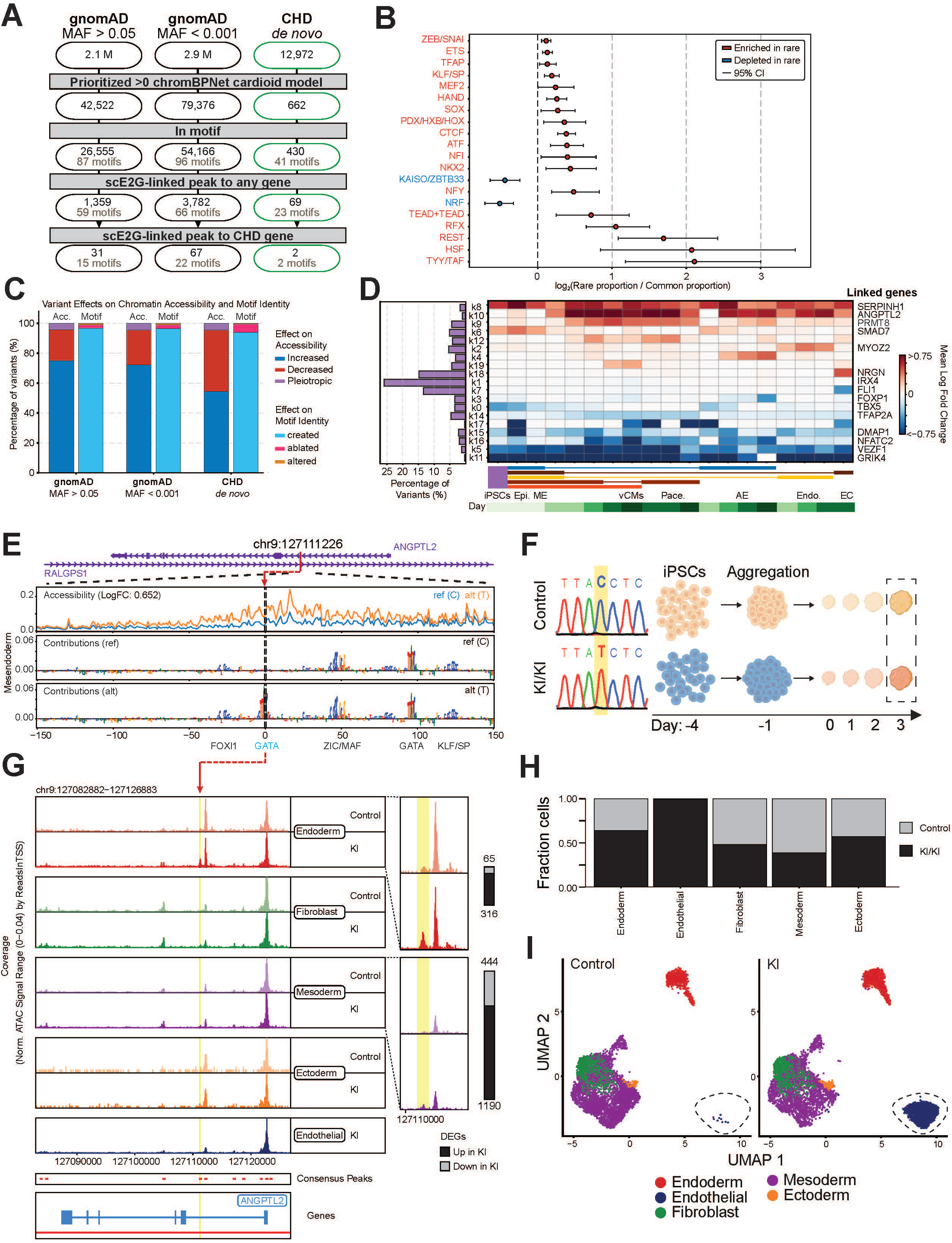
CHD-Derived Variants Disrupt Gene Regulation. (A) Prioritization pipeline identifies 100 rare noncoding variants linked to CHD genes from an initial pool of >5 million variants. (B) Rare gnomAD variants are enriched in cardioid transcription factor motifs compared to common variants. (C) Variants remodel regulatory landscapes through diverse motif alterations including creation, disruption, and bidirectional changes (D) CHD-derived noncoding variants exhibit cell state-specific effects with endothelial cells as the most disrupted lineage (E) *ANGPTL2*-linked variant (chr9:127111226:C→T) induces a gain-of-accessibility effect by creating a GATA motif. (F) Introducing chr9:127111226:C→T in the healthy male WTC11 cell line, scMultiomedata was collected at day 3 of cardioid differentiation. (G) *ANGPTL2* locus in the different clusters in Control and KI/KI shows a change of accessibility at the variant location (red/yellow highlights in the left/right panels, respectively), the right panel shows a zoomed-in scale. (H) Fraction of cells in each cluster in the Control and KI/KI samples. (I) Single-cell RNA analysis shows increased number of endothelial cells.

To test whether the specific motifs impacted by genetic variation show signatures of functional constraint, we compared the distribution of motifs hit by rare variants (MAF < 0.001) to those impacted by common variants (MAF > 0.05), using a multinomial enrichment analysis (**Methods**). Motifs disproportionately disrupted by rare variants included established cardiac regulators such as NKX2, HAND, and MEF2 as well as the TEAD+TEAD tandem homotypic configuration (**Fig. 7B)**. Among the prioritized rare variants, we identified four rare SNVs predicted to regulate *TBX5*, including rs377307764, which disrupts a GATA motif in a putative *TBX5* CRE (**Fig. S7A**) previously associated with CHD^96^. This variant is predicted to reduce accessibility beginning in the mesendoderm state, prior to detectable *TBX5* expression, while additional proximal variants show no measurable effect (**Fig. S7B**). We additionally identified 93 rare SNVs predicted to alter regulation of known CHD genes such as *NODAL, NOTCH1, NKX2-5* via disruption of GATA, CTCF and ETS family motifs.

A principal strength of our computational framework is its ability to predict directional effects on accessibility, enabling quantification of transcription factor-mediated tuning of chromatin accessibility. While the fraction of pleiotropic variants was similar across variant categories, the fraction of variants that decreased accessibility was greater in *de novo* CHD-derived variants (**Fig 7C**). Most prioritized variants are predicted to create a *de novo* motif that increases accessibility. In contrast, variants creating *de novo* binding sites for repressive transcription factors, such as *ZEB*/*SNAI* family members, were predicted to reduce chromatin accessibility (**Fig 7C**).

To prioritize noncoding variants with the highest likelihood of having a functional consequence, we focused on the subset residing within scE2G-linked regulatory peaks. Approximately 10% (5,368/54,595) lie within an scE2G-linked peak (**Table S17**). Pleiotropic effects were predicted for 177 rare/*de novo* variants within linked peaks, including variants that link to genes with known roles in heart development that aren’t broadly associated with CHD including *PAX2, GRHL2* and *ERBB3* (**Table S17**). We also identified a *TRIM16*-linked variant that creates a nested GATA motif within an OCT:SOX motif, which is predicted to reduce accessibility in pluripotency but increase accessibility in all somatic-like states (**Fig. S7C**), highlighting how complex motifs could influence multiple differentiation events across various lineages. Together, these findings support a model in which rare inherited noncoding variants contribute to CHD through complex, potentially oligogenic mechanisms.

To define the cell states disrupted by CHD-derived *de novo* variants^9^, we next evaluated the trends in predicted change in accessibility across ChromBPNet models. We observed that 62% (n=269) of variants that impact motif syntax impact multiple cell states (**Table S17**). Of the 161 variants that disrupt a single state, 83 (52%) were prioritized by the endothelial cell model while 9 (6%) were prioritized solely in the cardiomyocyte model (**Table S17**). A K-means clustering analysis demonstrated that the predominant category of variants comprised those predicted to confer modest alterations in chromatin accessibility across multiple cell states (n=108, **Fig. 7D**). However, the endothelial state was most disrupted when the two direction of effect categories were combined (*k*7+18, n=118, **Fig 7D**). Motif-associated variants within scE2G-linked peaks were predicted to regulate various genes implicated in cardiovascular development and function such as *TBX5*^97^, *FLI1*^98^, *IRX4*^99^, *TFAP2A*^100^ and *FOXP1*^101^. We additionally identified variants predicted to regulate genes with known roles in CHD-related, heritable outcomes, such as variants linked to *NTMT1* and *NFACT2* which are both associated with heart failure^102,103^ and a variant linked to *NAV1* which is critical during periods of brain development implicated in neurodevelopmental disorders, a frequent CHD comorbidity^104^. We additionally identified two unique variants associated with *SMAD7* regulation, a critical regulator of cardiogenesis^105^. These results highlight the need to expand variant validation efforts beyond cardiomyocytes and may explain the paucity of experimentally validated noncoding variants currently associated with CHD pathogenesis.

We additionally uncovered 202 variants that create known motifs de novo, 36 of which were within scE2G-linked-peaks (**Table S17**). One of these, an intronic SNV in *ANGPTL2*, located within a putative early endoderm cdCRE whose accessibility is driven by FOX, GATA, and KLF/SP motifs - is predicted to create a *de novo* GATA motif and increase chromatin accessibility in the mesendoderm and derived states (**Fig. 7D, E**). To test this prediction, we inserted this variant into a healthy hiPSC line, differentiated both control and knock-in lines into cardioids and generated scMultiomes (**Fig. 7F**). In agreement with the predicted creation of the GATA motif, we observed that this variant was sufficient to increase the containing peak’s accessibility (**Fig. 7G**) and to increase the number of endothelial cells within the cardioid (**Fig. 7H, I and Fig. S7D,E**). Ectopic *ANGPTL2* in adult tissue is associated with chronic inflammation in multiple organs, coronary artery disease and heart failure^106,107^. Ectopic expression was also noted in valve tissue from individuals diagnosed with aortic valve stenosis^108^. Its expression is also required for aortic valve development^109^ and vascular specification from the hemogenic endothelium^110^. However, it has not previously been associated with CHD, suggesting our pipeline can identify novel disease genes. Across all prioritized variants that impact motifs and link to genes, we nominate 154 candidate disease genes not previously associated with CHD. This result highlights how integrating scMultiome data with deep learning enabled noncoding variant interpretation can uncover disease-relevant genetic mechanisms.

## Discussion

In this study, we generated a high-resolution, scMultiome atlas of human cardioid differentiation. We leveraged this resource to train base-pair-resolution deep learning models that decode the cis-regulatory syntax governing cell fate and enable counterfactual prediction of cell context-specific effects of regulatory variants. Our models uncovered a latent layer of regulation encoded by predictive cell-state-specific transcription factor motif syntax, a feature obscured by conventional modeling approaches, that we employed to nominate novel disease variants, genes and regulatory mechanisms.

While our deep learning models learn a rich representation of the sequence features driving cell-state-specific chromatin accessibility, extracting biological insight requires a layered interpretation framework: nucleotide-resolution contribution scores to identify predictive sequence features, *de novo* motif discovery to define a unified regulatory lexicon, genome-wide predictive motif instance mapping, and *in silico* marginalization to quantify the isolated and combinatorial chromatin activity of individual motifs. Together, these methods enable analyses beyond conventional motif scanning to disentangle how combinations of TF motifs, often embedded in close proximity, synergize to drive cell state transitions. This model interpretation framework enabled us to assess the dynamic, lineage-specific activities of key regulators such as MEF2, to delineate the syntax of cell-state-specific regulation within cdCREs at loci like DHRS3, which is broadly required for retinoic acid signaling. We further show that TEAD activity is modulated by partner identity cooperating with SRF in cardiomyocytes and GRHL in ectodermal tissues and by the precise base-pair spacing constraints between the motifs. Together, these results advance our ability to interpret how TF motif combinations encode regulatory instructions in the noncoding genome and drive cell fate specification.

A second key capability of our framework is the prediction and mechanistic interpretation of variant effects on chromatin accessibility in a cell-state-specific manner. By coupling variant effect predictions with nucleotide-resolution attribution and motif disruption analysis, we can not only identify which variants alter accessibility but also define the precise regulatory features they disrupt. Critically, the diversity of early developmental cell states captured by our cardioid system spanning post-gastrulation transitions that are inaccessible to conventional genomic studies enables variant interpretation at the precise stages when cardiac lineages are first established. Applying this framework to CHD-derived variants, we identified 98 motifs affected by human genetic variants that exhibit either unidirectional effects on single cell states or pleiotropic effects across multiple states with respect to chromatin accessibility. We also identified many variants predicted to perturb regulation of genes not previously associated with CHD through cohort genetic studies, yet many have known roles in heart development and disease. De novo motif generation was predicted for 2,489 variants that connect to 2,641 genes, including a GATA motif within a CRE that targets *ANGPTL2*. Experimental validation of this variant provides mechanistic insight into the precise developmental contexts wherein noncoding variants may interfere with normal morphogenesis, culminating in CHD pathogenesis. Moreover, this approach establishes a framework for disease gene discovery and holistic quantification of genetic burden in diseases exhibiting oligogenic inheritance patterns as noncoding variants with low effect likely interact with additional, latent high effect variants.

Our findings additionally support the hypothesis that noncoding variants within the regulatory landscape of known disease genes can affect only a discrete subset of cell states, producing pathogenic mechanisms that are distinct from - and may not resemble - those caused by coding variants in the same gene. Because individual CREs can regulate multiple genes, a single noncoding variant may perturb several targets simultaneously, amplifying its pathogenic impact. Indeed 40% of prioritized, scE2G-linked CHD-derived variants are predicted to regulate 2 or more genes. These observations underscore the need for experimental model systems that faithfully recapitulate early development cell states to accurately interpret the functional consequences of genetic variation influencing developmental disorders such as CHD.

In summary, our study shows that it is possible to transition from descriptive atlases of human organogenesis to predictive, mechanistically interpretable models of the underlying cell-state resolved gene regulatory code. The deep learning and interpretation framework we present can serve as an *in silico* screening platform to systematically prioritize noncoding variants that disrupt cis-regulatory syntax across the precise developmental contexts in which cardiac lineages are established providing a direct path from genetic association to testable mechanistic hypotheses about congenital disease etiology. As the scale and resolution of single-cell genomic atlases continue to grow, frameworks of this kind will become increasingly powerful tools for decoding how noncoding variation disrupts the regulatory grammar of development and contributes to the full spectrum of congenital disorders.

### Limitations of our study

While our cardioid model successfully recapitulates early developmental events, its present cell state composition presents limitations for modeling the later, more complex aspects of organogenesis and disease. The current system lacks key supportive cell types known to be critical *in vivo*, such as neural crest cells. The absence of these lineages and their associated paracrine signals likely limits the maturation potential of the cell types that are present. Therefore, while our framework is powerful for dissecting early cell fate decisions, future studies aimed at modeling complex disease phenotypes will require more comprehensive organoid systems. Incorporating additional growth factors and mechanical cues, such as sheer stress, will be a critical next step in generating a more complete model of the human heart.

A second important limitation is that our variant interpretation framework is currently restricted to predicting effects on local chromatin accessibility. CHD pathogenesis can, however, involve variants that operate through a broad range of other molecular mechanisms including disruption of histone modifications, transcription initiation, elongation, splicing, post-transcriptional regulation, translation, and post-translational protein function. Variants acting exclusively through these mechanisms will not be captured by our models, and the fraction of CHD heritability attributable to such variants remains unknown. Encouragingly, predictive sequence-to-function models capable of predicting effects on diverse molecular readouts are rapidly expanding the scope of in silico variant interpretation^32,111–114^. Realizing the full potential of these models for disease variant prioritization will, however, require systematic collection of the relevant molecular data modalities in disease-relevant cell states and developmental contexts of the kind generated here.

## Methods

### 1. Cell culture

#### 1.1. Cell lines

The data was generated using two healthy male hiPSC lines^115,116^. CRISPR-interference experiments were conducted in an MYL7-GFP reporter line that encodes the doxycycline inducible CRISPRi machinery^117^.

#### 1.2. hiPSC cell culture

hiPSCs were cultured in mTeSR Plus medium (StemCell Technologies, USA, 100-0276) and on vitronectin (Thermo Fisher Scientific, USA, A31804). Cells were passaged every 3-4 days via dissociation with accutase (StemCell Technologies, USA, 07920) and cultured in ROCKi (Tocris Biosence, USA, 1254) for the first 24 hours-post-split.

#### 1.3. ANGPTL2 knock-in generation

The ANGPTL2-associated variant was inserted into the endogenous locus in a healthy male hiPSC line (WTC11^116^) using the sgRNA sequence 5’-GCAAAGATTGAAAGAGGTAA-3’ and single-stranded oligodeoxynucleotide donor sequence 5’-AGGGTCTAACTGAGAAGGTGGTATTTCAGCAAAGATTGAAAGAGATAAGGGAACAAACAACGAAGAAGGGGGCAGGGACCAGCAAGTATA-3’. Electroporation of the ribonucleotide protein complex was performed with the Neon Electroporation System (Thermo Fisher Scientific, USA) with 10 µL Neon tips and the following settings: Pulse V: 1300, pulse width ms: 30, pulse number: 1, 6µg SpCas9, 3.2µg sgRNA and 50pmol donor. hiPSCs were incubated with ROCKi for 1 hour pre-electroporation and 24 hours post in CloneR2 (StemCell Technologies, USA, 100-0691). Cells were - detached, electroporated, replated and allowed to recover for 5 days. Individual colonies were then picked, expanded and genotyped using the following primers: 239F, 5’-AGTCAAACGGGGTCACTCTT-3’ and 240R, 5’-GACCAACACAAACACCTCCC-3’. Twenty five independent colonies were picked and two colonies each were combined to generate the knock-in and control cardioids.

#### 1.4. Cardioid generation

The cardioid generation protocol was adapted from^34,35^. hiPSCs were seeded into 96-well plates at a density of 10,000 cells per well after achieving 70% confluency and dissociating with Accutase. Cells were resuspended in mTesR Plus medium with ROCKi and incubated overnight. On day 0, media was changed to FlyAB(Ins), containing RPMI (Thermo Fisher Scientific, USA, 11875119), B-27 with insulin supplement (Thermo Fisher Scientific, USA, 17504044), FGF2 (Fisher Scientific, USA, 23-3FB), BMP4 (Fisher Scientific, USA, 314-BP), LY294002 (Fisher Scientific, USA, 1130), CHIR (Tocris Bioscience, USA, 4423), and Activin A (R&D Systems, USA, 338-AC) in specified concentrations. Subsequent media changes were performed as follows: on day 1, with a refresh of FlyAB(Ins); after 36-40 hours, media was changed to BWIIFRa, which includes RPMI, B-27 with insulin supplement, BMP4, FGF2, IWP2 (Tocris Bioscience, USA, 3533), and retinoic acid (Tocris Bioscience, USA, 0695). Media was refreshed every 24 hours for days 2-4 with BWIIFRa. On days 5/6, media was changed to BFI containing RPMI, B-27 with insulin supplement, BMP4, and FGF2. On day 7, the media was switched to maintenance media with RPMI and B-27, refreshed every 48 hours.

#### 1.5. Cardioid dissociation and cryopreservation

At every time point (Days 0, 1, 2, 3, 5, 10), cardioids were dissociated into single cells with >90% live cells before freezing. TrypLE 10x (Gibco™, Thermo Fisher Scientific, USA, A1217701) was warmed to 37°C, and the centrifuge was cooled to 4°C. Cardioids were washed with DPBS on a bi-directional 37 µm filter (StemCell Technologies, USA, 27215) and then dissociated using prewarmed TrypLE 10x in a thermomixer set at 1000 RPM at 37°C for 20 minutes with gentle pipetting every 5 minutes. When cardioids were fully dissociated, cells were centrifuged at 300g for 5 minutes at 4°C, resuspended in PBS, and viability checks were performed, aiming for >95% live cells. Cells were thoroughly resuspended in a freezing medium (Bambanker, NIPPON Genetics, USA, BB05) and cryopreserved using a slow rate-controlled cooling process. Cryovials were then transferred to liquid nitrogen for long-term storage.

#### 1.6. Cardioid Thawing

Cryopreserved dissociated cardioids were transferred from liquid nitrogen to a bead bath until approximately half-thawed. The cell suspension was then immediately mixed with 1 mL of cold RPMI and transferred to a low-bind Eppendorf tube. The cells underwent two subsequent wash steps: they were centrifuged at 300g for 5 minutes at 4°C, and then gently resuspended in 1 mL of 1X PBS with 0.04% BSA (Thermo Fisher Scientific, USA, 15260037) using a wide-bore tip. After the final wash, the cell pellet was resuspended in an appropriate volume of 1X PBS with 0.04% BSA. To remove any remaining aggregates or debris, the final cell suspension was passed through a 70 µm cell strainer. The recovered cells were kept on ice, and their viability was assessed via trypan blue staining, with a quality control goal of greater than 95% viability.

### 2. Immunofluorescence and microscopy

Cardioids were washed with 1X PBS, fixed in 4% paraformaldehyde (PFA) for 20mins, cryoprotected overnight in 30% sucrose at 4°C, and embedded in Optimal Cutting Temperature compound (Sakura Finetek, USA, 4583). Samples were cryosectioned at 10 µm using a cryomicrotome (Thermo Scientific, USA, Microm HM 550), and adjacent sections were collected on coated glass slides (VWR, USA, VWRU48311-703). For immunofluorescence staining, sections were fixed again in 4% PFA for 20 minutes and washed with PBS. Samples were blocked with 5% goat serum (Abcam, USA, ab7481) in PBS containing 0.1% Triton X-100 (Sigma-Aldrich, USA, X-100) for 1 hour at room temperature, then incubated overnight at 4°C with primary antibodies (Table S18). After three PBS washes, sections were incubated with fluorescently conjugated secondary antibodies for 2 hours at room temperature and washed again. Finally, slides were mounted with Fluoromount-G (Southern Biotech, USA, 0100-01) and imaged using a Zeiss LSM 990 confocal microscope (Carl Zeiss, Germany).

### 3. Single-cell processing

#### 3.1. Single-Nucleus 10x Multiome Profiling

##### 3.1.1. snRNA-Seq Analysis

Single-nucleus libraries were generated using the 10x Genomics Chromium scMultiome ATAC + Gene Expression system, sequenced on an Illumina platform, and processed via Cell Ranger ARC. Downstream single-cell RNA (scRNA) analysis was performed in Seurat (v5.0). To strictly control for background contamination, ambient RNA signatures were estimated and computationally subtracted using the decontX algorithm^37^. Following the removal of low-quality cells, RNA counts were normalized utilizing SCTransform (v2) and subjected to PCA and UMAP dimensionality reduction. To specifically interrogate developmental heterogeneity, the dataset was clustered using consensus variable features, and manually annotated into specific lineages based on canonical marker expression. Differential gene expression between Knock-In (KI) and control genotypes was assessed within each cell type using a Wilcoxon rank-sum test.

##### 3.1.2. snATAC-Seq Analysis

Analysis of single-nucleus ATAC-seq (snATAC) data was performed using ArchR^118^. Nuclei were strictly filtered for Transcription Start Site (TSS) enrichment (≥ 5), unique fragments (≥ 1,000), and singlet status, followed by the enforcement of a joint multiomic whitelist to retain only high-quality cells present in both modalities. Dimensionality reduction was executed via Iterative Latent Semantic Indexing (iLSI), and RNA-derived cluster annotations were computationally transferred to the ATAC-seq embeddings. To capture genotype- and lineage-specific regulatory elements, reproducible peak sets were called using MACS2^119^ on pseudo-bulk grouped cells.

#### 3.2. SHARE-seq

SHARE-seq protocol was adopted from ^120,121^, and a detailed protocol is provided in Note S1.

#### 3.3. Library Sequencing

All DNA libraries were sequenced on a NextSeq 500/550 High Output Kit v2.5 (150 Cycles, 20024907), NovaSeq X Series 25B Reagent Kit (300 cycles, 20104706), NovaSeq 6000 S4 Reagent Kit v1.5 (300 cycles, 20028312) with XP workflow. Paired-end sequencing was run with a 96-99-8-96 configuration (Read1-Index1-Index2-Read2). We quantified DNA libraries using Qubit and tapestation, then prepared library pools at 4nM concentration. Sequencing was performed at the Stanford Genome Technology Center (NovaSeq6000), NovoGen (NovaSeqX) and in-house instrument (NextSeq550).

### 4. SHARE-seq data preprocessing

Raw sequencing data was preprocessed using a customized pipeline suited to SHARE-seq https://github.com/GreenleafLab/shareseq-pipeline (stable release v1.0.0) ^122^. Fastq files were de-identified prior to public deposit using BAMboozle^123^.

### 5. Data analysis

A brief description of the analysis steps is provided in the following sections.

#### 5.1. SHARE-seq data QC and filtering

We performed per-sample QC filtering by manually inspecting and thresholding the following metrics: 1) TSS enrichment ratio and number of fragments for ATAC fragment files, 2) number of UMIs, number of genes and percentage of mitochondrial reads for RNA sparse matrices, 3) ratio of RNA UMIs vs ATAC fragments to remove cells with low quality in one modality. All sample filtering thresholds are summarized in Table S2.

#### 5.2. RNA normalization, ambient RNA removal, dimensionality reduction, and clustering

We used Seurat (v5.0)^36^ in R (v4.2.2) to process filtered RNA sparse matrices into Seurat objects. We adopted an iterative dimensionality reduction and clustering workflow to sequentially annotate cell types. For each iteration, we first performed SCTransform v2 and variable feature selection on RNA raw counts of each sample, then selected the top 3,000 consensus variable features across samples using the SelectIntegrationFeatures function from Seurat, excluding mitochondrial genes, sex chromosome genes, and cell cycle genes to minimize batch effects. We merged the raw RNA counts from per-sample objects into a single matrix, performed SCTransform v2 using consensus features, and used the DecontX function from the celda (v1.6.1) package^37^ to remove ambient RNA contamination per cell. The decontaminated counts were then scaled to 10,000 UMIs per cell and log-normalized. Similar to the process mentioned above, we selected a list of the top 3,000 consensus variable features. PCA was performed on the merged object with the consensus features, followed by cell clustering using the Louvain algorithm at a resolution of 0.3 with 50 PCs and UMAP embedding. We then inspected each cluster and removed any low-quality clusters with significantly lower UMIs than other clusters. After removing cells in the low-quality clusters, we repeated the processing steps starting from RNA raw counts for each sample. This process was repeated until no more low-quality clusters were identified, which required three iterations. Cells in the final set of clusters passing this iterative QC were considered “whitelisted”. All final cluster annotations are included in Table S3. All final retained cell barcode can be found in Table S5.

#### 5.3. Differential Gene Expression Analysis

All differential gene expression analyses were performed using the decontaminated counts stored in the decontX assay of the integrated Seurat object.

First, to identify cell-state-specific marker genes across the entire dataset (combining genotypes), the FindAllMarkers function was run using both the MAST (test.use = "MAST") and Wilcoxon rank-sum (test.use = "wilcox") tests. Both tests were executed with the following parameters: min.cells.feature = 20, min.pct = 0.1, min.diff.pct = 0.2, and logfc.threshold = 0.3. A consensus marker list was generated by retaining only the genes identified in the intersection of both test results. All marker genes per cell state can be found in Table S6.

#### 5.4. scATAC data processing and label transfer

We used ArchR (v1.0.2)^118^ to process filtered ATAC fragment files into a single ArchR project. After filtering to the final whitelisted cell barcodes from the iterative RNA processing workflow and transferring the clustering and cell type annotations, we called peaks per cluster using Macs2 (v2.2.7.1)^119^, merged peaks into a single reproducible peak set using ArchR’s iterative overlap strategy, and created a cell-by-peak matrix of fragment counts. We identified mcarker peaks per cluster using a Wilcoxon rank-sum test. For final cluster annotation, we use the well-defined RNA cluster labels and store them as metadata in the ArchR object.

#### 5.5. scE2G

We used the scE2G*^Multiome^* model to predict enhancer-gene connections in each cell type (https://github.com/EngreitzLab/scE2G/tree/v1.2)^63^. Briefly, scE2G first defines a set of candidate element-gene pairs where candidate elements are derived from peaks called on the pseudobulk ATAC-seq data from each cell type and are paired with genes with a transcription start site within 5 Mb. We used the model’s default TSS reference file with some manual adjustments to select the appropriate TSS for each gene. Next, it annotates each pair with seven features calculated from the multiome data, including 1) a pseudobulk Activity-By-Contact (ABC) score, where 3D contact is estimated by an inverse function of genomic distance; 2) the Kendall correlation across single cells between element accessibility and gene expression; 3) whether the gene is “ubiquitously expressed”, and 4) several other measures of genomic distance and chromatin accessibility around the element and promoter. Then, the features are integrated to assign each candidate element-gene pair a score representing the likelihood that the element regulates expression of the gene. Regulatory enhancer-gene interactions were defined as element-gene pairs with a score greater than 0.177, which is as the score yielding 70% recall when evaluating predictions in K562 cells against CRISPRi-validated enhancer-gene pairs^124^.

#### 5.6. Azimuth

To compare our *in vitro* cell states to an *in vivo* reference, we performed a projection analysis using Azimuth^36^ (v2.1.0). The decontaminated scRNA-seq counts matrix from the integrated cardioid dataset was used as the query. We ran the RunAzimuth function to map these query cells onto the human gastrula reference atlas ("hGas"). The mapping was performed using pre-computed reference data, and cell annotations were transferred at two levels: celltype.l1 (broad developmental lineages) and celltype.l2 (specific cell identities). The resulting object, containing the projected UMAP coordinates and cell predictions (predicted.celltype.l1, predicted.celltype.l2, and mapping.score), was saved for downstream analysis. The metadata, including query cluster labels and all predicted annotations and scores, are available in Table S8.

#### 5.7. Optimal transport

We applied Waddington-OT^53^ to model single-cell trajectories for each modality, with default pipeline settings. A cell at a time *t* is represented by the gene-expression vector *x* ∈ *R^G^*, where *G* is the number of genes. For each adjacent pair of time points (*t_i_*, *t_i_*_+1_), we first carried out a pairwise principal-component analysis, retaining the top *d* components shared by both time points. Coordinates in this *R^d^* subspace served as input to an entropic, unbalanced optimal-transport problem ^125^. Only the row sums were constrained, allowing the column sums to vary so that proliferation or death could be accommodated. Solving this optimisation yielded a coupling matrix 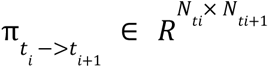 Matrix multiplication of the cell embedding matrix at *t_i_* with 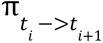 denotes the mass transport between cells in *t_i_* to *t_*i*_*_+1_. Assuming Markovian dynamics, long-range couplings were approximated by successive matrix multiplication: 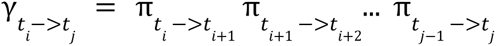. For each starting cell, the probability mass in γ arriving in user-defined final clusters was summed to create an aggregate fate matrix *F* ∈ *R*^*N*×*K*^, where *N* is the total number of cells and K is the number of fates. The full pipeline was executed three times—once on RNA expression, once on ATAC gene scores, and once on a concatenated RNA + ATAC feature matrix. Within each modality, we trained separate models on control cells, on knockdown cells, and on the combined dataset, enabling direct comparison of transport maps and fate probabilities across perturbations. Final differentiation trajectories by gene expression and OT analysis can be found in Table S7.

#### 5.8. ChromBPNet

##### 5.8.1. Training

For each cardioid cell state, ATAC-seq fragments from cells constituting that cell state were aggregated. These aggregated fragments were fed into the ENCODE peak calling pipeline to identify peaks in each cell state. The per-cell state aggregated fragments and peaks were then used to train ChromBPNet (https://pubmed.ncbi.nlm.nih.gov/39829783/) models.

The ChromBPNet models were trained using the framework provided in the ChromBPNet Python package (https://github.com/kundajelab/chrombpnet). For each cell state, 5 individual ChromBPNet models were trained by following the 5-fold cross-validation scheme used in the ChromBPNet package, in which each chromosome is present in the test set of at least one cross-validation fold. The chromosomes used per fold are shown in Table S9. All training was done using the hg38 reference genome. All of the ChromBPNet models were trained using a Tn5 bias model trained on the background regions from the fold 0 chromosomes of the vCM cell state. The per-fold background regions were defined by the ChromBPNet package. A bias threshold factor of 0.6 was used.

##### 5.8.2. Per-cell state predictions

For every cell state, once the 5-fold models were trained, predictions for that cell state were made by averaging individual predictions from the 5-fold models. The log counts prediction for the cell state was the average of the log counts predictions of all the individual folds. The profile logits prediction for the cell state was the average of the profile logits predictions of all the individual folds. The profile logits prediction for the cell state was then softmaxed to produce the profile prediction for the cell state. The log counts prediction for the cell state was exponentiated and then multiplied by the profile prediction for the cell state to produce the counts-scaled profile prediction for the cell state. This averaging procedure was used for both the with-bias and de-biased model predictions.

##### 5.8.3. Generating contribution scores

We used the DeepSHAP^58^ implementation of the DeepLIFT algorithm^56^ to compute the contribution that each base has towards a model’s prediction for the total log counts over a genomic sequence. The contribution scores in each cell state are computed as the average of the contribution scores from each of the 5-fold models. Contribution scores were calculated on the de-biased models.

##### 5.8.4. Identifying a motif lexicon

In each cell state, we identified the motifs of transcription factors that are active in that cell state and contribute towards chromatin accessibility predictions. To do this, for each cell state, we began by computing contribution scores for all 2114 bp sequences centered at every peak summit identified in that cell state. Then, TF-MoDISco ^57^ (https://github.com/kundajelab/tfmodisco) was run on the per-cell-state contribution scores to identify motifs present in that cell state. The *modisco motifs* command was run with 1,000,000 seqlets (the *-n 1000000* option).

Once motifs were identified in each cell state, the motifs were clustered to create a non-redundant set of motifs. The 2169 motifs identified across all 59 control and knockdown cell states were clustered using the MoitfCompendium^59^ (https://github.com/kundajelab/MotifCompendium) Python package to yield a set of 418 distinct motifs. Clustering was done using default clustering options at the 0.95 similarity threshold. The 418 distinct motifs were further classified into 172 motif groups by automated annotation (MoitfCompendium), followed by manual curation. The 172 groups were next clustered into 106 distinct motif families (Note S3).

##### 5.8.5. Marginal footprinting

For each motif and for each cell state, an *in silico* marginal footprint score was computed to quantify the effect that motif has in that cell state. First, for each model, 1000 of the training background sequences identified during ChromBPNet model training were sampled. Then, the predicted counts from the de-biased model were computed (in this case, the predicted counts were computed as the average of the predicted counts of the forward and reverse sequence). This served as a baseline for the amount of accessibility latent in these background regions. Then, for each motif, a consensus DNA sequence was determined by taking the maximum contributing base pair at every position. For each motif, that motif’s consensus sequence was then inserted into the center of the 1000 background sequences. Predictions from the de-biased models were made on these sequences with *in silico* motif insertions in them (the motifs were inserted at the center of the sequence by replacing the bases that were already there); the predicted counts of the forward and reverse sequences were averaged to produce the predicted counts of these sequences. The log2 fold change between the predicted counts of the motif-inserted-sequence and the predicted counts of the without-motif-sequence was then computed per sequence. This log2 fold change was then averaged across all 1000 sequences to produce an effect size for the marginal effect of the motif in that model. These effect sizes were then averaged across the five-fold models to give cell-state-specific marginal footprint scores of the motif. This process was done per motif to give marginal footprint scores for each motif in each footprint.

These marginal footprint scores were plotted as a line plot across the differentiation trajectory to produce Fig. S3H and Note S3. These marginal footprint scores were row-normalized and then plotted in a heatmap to produce Figure 3F-H. Figure 3H shows rows of Figure 3G, which have been subset to highlight differences in the marginal footprint effect sizes of different motifs in the same family.

##### 5.8.6. Annotating motifs in each cell state’s regulatory landscape

We performed a comprehensive identification of motifs present in the regulatory regions of every cell state. For each cell state, we used the FiNeMo (https://github.com/kundajelab/Fi-NeMo) Python package to identify all occurrences of each motif in the non-redundant motif set within the summit-centered peak sequences of that cell state. The result was an annotation of all motif instances of every motif within the peak sequences of every cell state. The *finemo call-hits* command was run with default options.

##### 5.8.7. Motif enrichment in peaks

We computed motif enrichment within cell states across peak clusters (Figure S3C). To do this, we first began by selecting two peak clusters of interest in a cell state. Then, we counted the number of motif instances of each type of motif present in that cell state in each peak in the two peak clusters. Then, for each motif, we created a contingency table where the top left was the number of instances of that motif in all peaks in the first peak cluster, the top right was the number of motif instances of that motif in all peaks in the second peak cluster, the bottom left was the total number of other motif instances in all peaks in the first peak cluster, and the bottom right was the total number of other motif instances in all peaks in the second peak cluster. Then, an odds ratio and p-value were computed for each contingency table using Fisher’s exact test.

##### 5.8.8. Gene-motif linking analysis

The first step of the gene-motif linking analysis was to filter genes. First, for each gene, the maximum expression of the gene across all cell states in normalized units of logTPM was computed. Then, any gene whose maximum expression is < 1% of the maximum expression (in units of logTPM) of any gene in any cell state is removed.

Then, the next step is to identify linked peaks for each gene. For each gene and each cell state, all scE2G links to that gene in that cell state were considered. If any of those links overlapped a peak in that cell state, that peak was added to the total set of peaks linked to that gene in that cell state. This was done across all cell states to yield a comprehensive set of scE2G-linked peak regions that are linked to that gene per cell state.

Then, for each cell state, gene, and motif, all instances of that motif identified in peaks linked to that gene in that cell state were identified as *gene-linked motif instances*. For each one of those gene-linked motif instances, the sum of the contribution scores of that motif CWM was scaled based on its distance to the gene (the distance weighting function is *e^-(distanceToTSS/5000)+1/e*) to yield a *distance-weighted motif to gene contribution score*. The distance-weighted contribution scores of all instances of a motif were summed to compute a score for how much that motif contributes to that gene in that cell state. This was done across all combinations of motifs, genes, and cell states.

#### 5.9. Consensus non-overlapping peakset

A consensus, non-overlapping peakset was constructed by merging peaks from every cell state. For each cell state, each peak set was transformed into a list of 500 bp bins centered around each peak summit. Then, these summit-centered bin lists were combined across all control and knockdown cell states, leading to a list of 9,643,338 500 bp regions. The peak calling pipeline, which uses macs3, annotates each summit with a p-value indicating how likely that summit is really a peak summit. The combined summit-centered bin list was then sorted by macs3 p-value, with the bins with the lowest p-value (most likely to be a peak summit) at the top and bins with the highest p-value (least likely to be a peak summit) at the bottom. At this point, the consensus peakset is an empty set and contains no summit-centered bins. Then, each summit-centered bin in the sorted list was considered, and the non-overlapping peakset was built with the following rule: if the bin had any overlap with any bin already in the peakset, the bin was not retrained in the final peakset; however, if the bin did not overlap with any bin that was already in the peakset, it was added to the peakset. Through this procedure, a non-overlapping summit-centered peakset of 628,378 500 bp bins was created.

#### 5.10. Peak clusters

For each of the 628,378 peaks in the consensus peakset, its total accessibility is computed per cell state as the sum of the total number of Tn5 insertions in the peak region in that cell state. This gives a peak by cell state counts matrix. This raw counts matrix is then normalized by the read depth in each cell state so that each element in the matrix, which corresponds to a specific peak in a specific cell state, is divided by the read depth in that cell state. Then, a copy of the normalized counts matrix is made which is then column-wise z-score normalized. Within each column (corresponding to a specific peak), this z-score normalization is done by first computing the column-wise mean (the mean normalized accessibility across all cell states) and then subtracting this mean from all values in the column, and then computing the column-wise standard deviation (the standard deviation of normalized accessibility across cell states) and then dividing the column values by this standard deviation. Then, the normalized counts matrix and the z-score-normalized counts matrix are concatenated along the cell state axis so that each peak is now represented by a vector of the normalized counts value across all cell states, concatenated with the z-score-normalized values across all cell states. Then, the peaks are clustered using scikit-learn’s Mini-Batch K-Means clustering with k-20 (*sklearn.cluster.MiniBatchKMeans(n_clusters=20, random_state=0, batch_size=1000, n_init=50)*). Three pairs of clusters with similar dynamics were merged (original clusters 6 and 16 were merged, original clusters 1 and 11 were merged, and original clusters 8 and 19 were merged). Finally, the clusters were manually re-ordered. This led to 17 peak clusters.

#### 5.11. Motif changes in knockdown

Every control cell state has a corresponding knockdown cell state. For every control/knockdown cell state pair, for each peak in the consensus peakset, we compute the log_2_ fold change between the normalized counts per million in the control and knockdown cell states; a positive log2 fold change indicates that the peak was more accessible in the control than the knockdown, and a negative value indicates that the peak was more accessible in the knockdown than the control. Peaks that had no counts in neither control nor knockdown were removed from this analysis for that cell state pair. Then, the remaining peaks were clustered by their log2 fold change value using k-means clustering with k=3. For each cell state pair, this separated the peaks into 3 clusters: a cluster containing peaks more accessible in control than knockdown, a cluster containing peaks more accessible in knockdown than control, and a cluster containing peaks with similar accessibility across both control and knockdown cell states.

Then, for each motif, we created a contingency table where the top left was the number of peaks in the control-accessible peakset that contained any instance of that motif family, the top right was the number of peaks in the knockdown-accessible peakset that contained any instances of that motif family, the bottom left was the number of peaks in the control-accessible peakset that did not contain any instances of the motif family, and the bottom right was the number of peaks in the knockdown-accessible peakset that did not contain any instances of the motif family. Then, an odds ratio and p-value were computed for each contingency table using Fisher’s exact test. The odds ratio and p-value of differential motif enrichment between control and knockdown are shown for each motif family and each control/knockdown cell state pair in Figure 5H.

#### 5.12. Integration of Data from two cell lines

Endothelial cells represented a rare population in the initial SHARE-seq experiment, yielding insufficient ATAC-seq fragments (3.5M) to train a robust ChromBPNet model. To address this limitation, an additional batch of cardioids was generated from the same two healthy male hiPSC lines and processed identically. ATAC-seq fragments from endothelial cells across both experimental batches were pooled, increasing the total fragment count to 12M. Peak calling was performed on the combined fragment set using the ENCODE peak calling pipeline, following the same procedure as described for other cell states. A Tn5 bias model was trained specifically for the endothelial cell fragments using a bias threshold factor of 0.5. This bias model served as the basis for training ChromBPNet models for endothelial cells, following the same 5-fold cross-validation framework and prediction procedures applied to all other cell states. This integrated dataset was used for motif discovery and variant scoring in Figures 6 and 7.

#### 5.13. Variant Effect Prediction with ChromBPNet

We made *in silico* variant effect predictions of noncoding variants using our ChromBPNet models. Each variant was specified by a genomic coordinate, a reference allele (which matched the hg38 reference genome), and an alternate allele. For each variant, we took the 2114 bp genomic sequence centered around the variant to be the “reference sequence”. We substituted the center reference allele for the alternate allele to create the “alternate sequence”. To predict the effect of a variant in a model, we compared the model’s prediction on the reference sequence to the prediction on the alternate sequence. All predictions were made using the de-biased models.

When comparing the reference and alternate predictions, a variety of scores are computed to quantify the effect of the variant. These include the log2 fold change (LFC) of the total predicted coverage (total counts) over the two sequences and the active allele quantile (AAQ), which is a measure of how the predicted counts in the reference and alternate sequences compare to the distribution of total counts in all peak regions used to train the models. More information about these scores and how they are computed can be found in the variant-scorer Python package (https://github.com/kundajelab/variant-scorer). To compute cell-state-level scores, the LFC and AAQ were averaged across the 5-fold models.

#### 5.14. Variant Datasets and Scoring

Three sets of noncoding variants (SNVs + indels) were scored using the ChromBPNet models described above. The first set consisted of 12,972 de novo variants from the Pediatric Cardiac Genomics Consortium (PCGC)^9^. The second set was derived from gnomAD v4.1^91^, downsampled and filtered to retain only variants with a maximal population allele frequency (grpmax) below 0.001 (n=2,890,669), representing a rare variant set enriched for potentially functional alleles. The third set consisted of common variants drawn from gnomAD (allele frequency > 0.05, n=2,110,172), used as a negative control set under the assumption that variants at high population frequency are less likely to disrupt critical regulatory elements. The gnomAD variant sets were randomly downsampled prior to scoring to reduce computational load; PCGC de novo variants were retained in full without downsampling (Figure 7A).

All variants were scored across all 29 ChromBPNet models trained on scATAC-seq fragments from cardioids as described above. For each variant in each model, the LFC of predicted accessibility, AAQ, and overlap with ATAC-seq peaks identified in that cell state were computed.

Variants were said to be prioritized in a given model if they met the following criteria: (1) predicted to significantly change accessibility (|LFC| > 0.25), (2) had an AAQ > 0.05, and (3) either overlapped an ATAC-seq peak called in that cell state, or were located outside of a peak but showed a significant positive LFC (potentially creating a *de novo* accessible region). Variants that were prioritized by at least one model were retained for all subsequent analyses.

Additionally, variants were assessed for their impact on transcription factor binding motifs: variants falling within a predicted existing motif in the reference sequence or predicted to create a new motif were flagged as "in motif." For variants flagged as "in motif," only those affecting motifs predicted to exist in the reference sequence were considered when requiring overlap with scE2G peaks, as variants predicted to create new motifs lack corresponding information about de novo peak formation.

#### 5.15. Motif enrichment analysis in rare versus common variants

For all prioritized variants, DeepSHAP contribution scores were computed for both the reference and alternate allele sequences. FiNeMo was then run on both the reference and alternate sequences to identify motif instances present in the 2114 bp window centered on each variant position at the cell state with the highest predicted effect.

To assess whether rare noncoding variants preferentially disrupt distinct regulatory motifs compared to common variants, we performed a multinomial-based enrichment analysis.

Prior to analysis, motifs matched with variants with fewer than 10 total observations across rare and common variant sets were excluded to reduce noise from sparse categories. Motif hit counts were aggregated across all cell states for each variant set. To assess motif enrichment, we established a null distribution through 100 iterations of split-half resampling of common variants, then compared rare variant motif distributions against this baseline using bootstrapped confidence intervals (1,000 iterations) to identify motifs exhibiting significant deviations between common and rare variant sets generating synthetic rare and common variant sets from the fitted multinomial distributions (**Fig. 7B**). Motifs were considered significantly enriched or depleted in rare variants if their 95% CI excluded zero. This approach assumes that common variants reflect neutral regulatory variation and that multinomial sampling adequately captures the distribution of motif hits across prioritized variants.

#### 5.16. Motif-level variant impact analysis

By comparing the motif annotations and contribution scores between reference and alternate sequences, each variant was classified according to its predicted mechanistic impact on transcription factor binding. A variant could: (1) have no detectable predicted effect on any motif, (2) be predicted to increase or decrease the accessibility of an existing motif binding site (motif modulation), (3) be predicted to abolish an existing motif binding site or create a de novo motif binding site (motif disruption/creation), or (4) be predicted to change a motif binding site into one that binds a different transcription factor (referred to here as pleiotropic effects).

#### 5.17. Gene-level variant linking

To connect variants to potential target genes, we leveraged the scE2G peak-to-gene links described above. For each prioritized variant, we identified all scE2G links that: (1) originated from a peak overlapping the variant position, and (2) were present in at least one cell state in which the variant was prioritized. This yielded a set of variant-gene pairs with predicted regulatory connections. We further refined this set by restricting to variants linked to known CHD genes.

## Data and Code accessibility

Available upon request.

## Acknowledgements

We thank Nick Addleman, John Coller, and the Stanford Genome Technology Center for their assistance with sequencing libraries, Timothy Nelson and the Todd and Karen Wanek Family program for healthy hiPSC lines, and members of the Gifford, Greenleaf and Kundaje labs for their helpful discussions. This work was supported by RM1HG007735, R01NS128028, R01HL171611, DP1HG013599, R01HG013317, UM1 HG012660, U01HG011762 (to S.B.M), UM1HG011972 (to W.J.G); support from the Stanford Cardiovascular Institute, Saving Tiny Hearts Society and Additional Ventures (to C.A.G); support from NHGRI Impact of Genomic Variation on Consortium (UM1HG011972); the Novo Nordisk Foundation Center for Genomic Mechanisms of Disease (NNF21SA0072102); NHLBI R01HL159176; the Applebaum Foundation; Gordon and Betty Moore; and the BASE Research Initiative at the Lucile Packard Children’s Hospital at Stanford University (to J.M.E). W.J.G. is an Arc Innovation

Investigator and Director of the Stanford RNA Medicine Program. C.A.G. is the Akiko Yamazaki and Jerry Yang Faculty Scholar in Pediatric Translational Medicine, Stanford Maternal & Child Health Research Institute and Director of the Mann Charities Biobank at the Betty Irene Moore Children’s Heart Center, Lucile Packard Children’s Hospital, Stanford. M.U.S. acknowledges the support of an NSF Graduate Research Fellowship (DGE-1656518) and a graduate fellowship award from Knight-Hennessy Scholars at Stanford University. M.A. acknowledges the support from the AHA postdoctoral fellowship. E.G.P acknowledges support of the *Training Grant in Myocardial Biology at Stanford (NIH 5T32HL094274-13)*. J.M.S acknowledges support of the National Science Foundation (NSF) for a Graduate Research Fellowship (DGE-2146755). E.M acknowledges the support from the HFSP postdoctoral fellowship. This research was made possible through the Allen Cell Collection, available from Coriell Institute for Medical Research. We’d also like to thank all the families affected by congenital heart disease that agree to be part of research studies.

## Author Contributions

E.M., R.Z. and S.D. contributed equally as co-first authors. E.M., W.J.G., and C.A.G. conceived the study; E.M., J.P., E.G.P., J.M.S and J.R. optimized cardioid cultures and performed sample collection, with input from C.A.G.; E.M. conducted SHARE-seq experiments; E.M. programmed and performed data preprocessing, with input from B.B.L. and S.H.K.; E.M., E.G.P. and M.M. annotated all cell types, with input from C.A.G.; R.Z. and S.D. trained ChromBPNet models and performed analyses related to the motif lexicon and transcription factor cooperativity, with input from A.K.; M.U.S performed scE2G processing and analysis, with input from J.M.E.; I.A. programmed and performed optimal transport analysis, with input from E.M.; M.A. and J.M.S performed immunofluorescence imaging, with input from C.A.G.; Y.Z., I.E., S.D. and C.A.G. performed causal variant analysis, with input from A.K.; E.M., R.Z., S.D., W.J.G., A.K., and C.A.G. wrote the manuscript, with input from all authors.; A.K., W.J.G., and C.A.G. jointly supervised the work.

---

**Figure S1.**
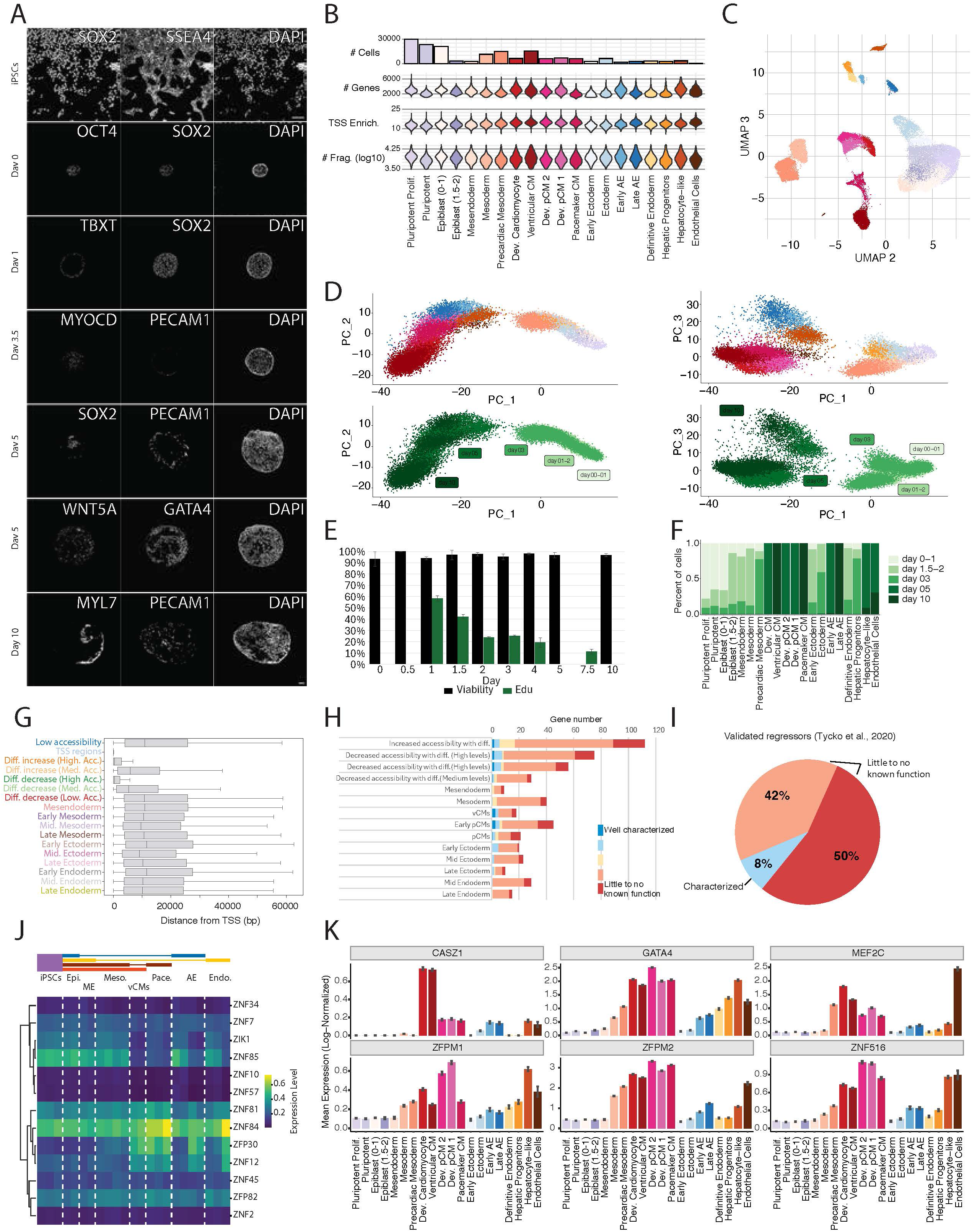
(A) Raw channel intensities. Pluripotency: *SOX2*, *SSEA4*, *OCT4*; Early mesendoderm specification: *TBXT*; Endothelial cells: *PECAM1*; Early ectoderm: *SOX2* on day 5; Mesoderm: *GATA4*; Ectoderm: *WNT5A*. (B) single-cell QCs per cell type. (C) UMAP 2 and UMAP 3 embeddings, corresponds to scRNA data in Fig. 1B. (D) PCA plots. Left - PC1 vs PC2 captures the general time component of cardioid differentiation. Right - PC3 captures the variance between the lineages. The top panels are annotated by cell type, and the bottom panels are annotated by time point. (E) Black bars: Viability of harvested cells during cardioid differentiation. Green bars: EdU staining. Proliferation drastically decreases in the first few days of differentiation. (F) The time point composition of each cell type shows that later time points are more prevalent in more mature cell types. (G) Distance of peaks in the peakset from the nearest TSS. (H) Known and predicted function of 502 ZNFs with cell-state specificity, raw numbers for Fig. 1F. (I) Fraction of known function for the 13 ZNFs with validated repressor domain. Out of the 13 ZNFs, only *ZNF57* was annotated as “Characterized”, mainly because of its known activity in methylation imprinting, also in embryonic development. (J) Log-normalized expression, corresponding to Fig. 1H. (K) Log-normalized expression across cell states.

**Figure S2.**
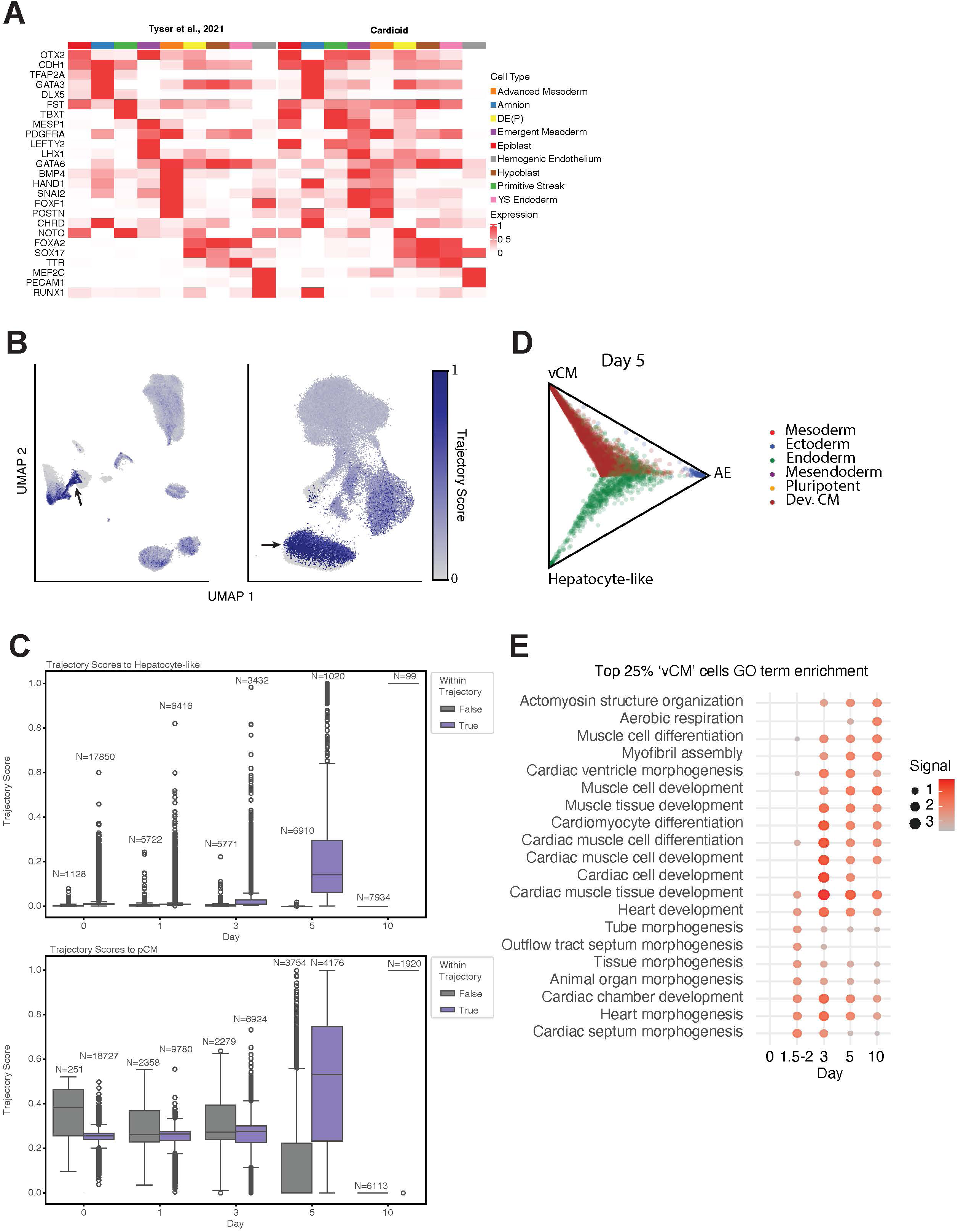
(A) Human gastrula cell type marker expression from Tysler et al.^3^, Extended Figure 2A and mapped cardioid cells gene expression. Left and right, respectively. (B) Optimal Transport Trajectory analysis identifies distinct cells predicted to differentiate to Pacemaker CM state (Black arrow). (C) Same as Fig. 2D but for Hepatocyte and Pacemaker CM trajectories. (D) The Optimal Transport Fate Matrix by Day displays the Fate probabilities for each cell at Day 5 of differentiation toward either of the final states (Late AE, vCMs, Hepatocytes). (E) GO term enrichments during cardioid development for the top 25% of cells with the highest probability of becoming vCMs.

**Figure S3.**
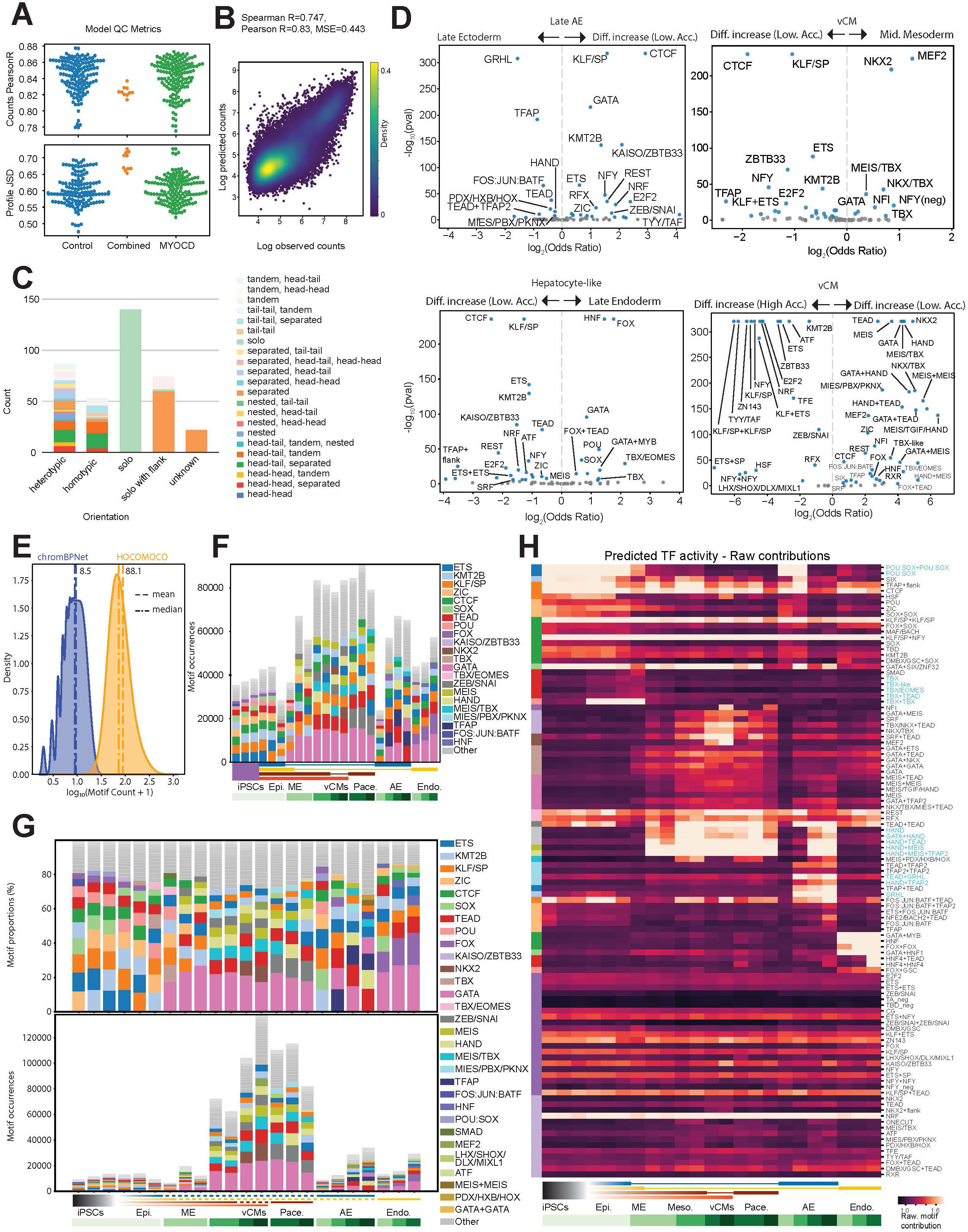
(A) ChromBPNet QC. Counts Pearson R of the control and MYOCD KD cardioids. For the Endothelial cells, we merged the fragments to train the model (“Combined”). All models achieved correlations ranging from 0.78 to 0.88, with a median of 0.85. We also quantified the extent to which the base pair-resolution predictions of the model were accurate by computing the Jensen-Shannon Distance (JSD) between the observed and predicted distributions of counts across bases per peak. For each cell-type-specific model, the median JSD among all peaks in that cell-type was an average of 0.61. (B) Correlation of predicted vs observed counts for the control vCM cluster. (C) Annotated subtypes of identified motifs in our data showed complex composition possibilities for homo/heterotypic motifs. (D) Log odds ratios of motif abundance between different peak sets in Fig. 3C. (E) Number of motifs per peak for ChromBPNet and HOCOMOCO V12 (blue with mean=8.5; median=8 and orange with mean=88.1; median=72, respectively) (F) Same as Fig. 3D but with all motifs. The top 10 abundant motifs in each cell state are annotated. (G) Motif proportion and occurrences for Fig. 3C, peak cluster “Mid. Mesoderm”. (H) Log_2_FC values for Fig. 3G. Highlighted motifs indicated by blue font.

**Figure S4.**
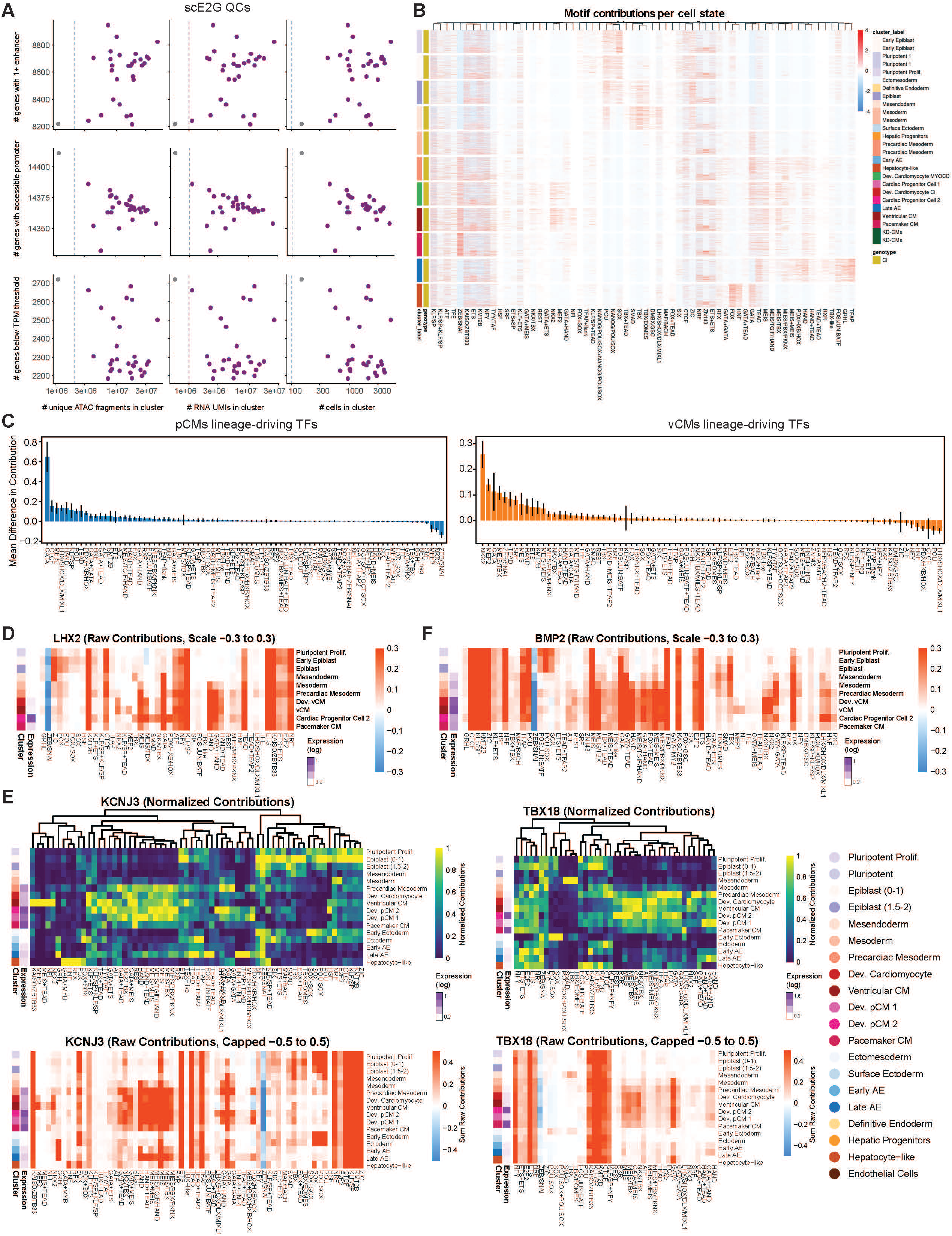
(A) scE2G QC. All cell states passed the QC threshold. (B) in genome contribution for the entire motif compendium across all cell states. The sums of contributions (intensity) for each motif family are column-scaled across all expressed genes (approximately 12,000 genes) in each cell state. (C) Average contributions for each motif in pCM and VCM progenitors (left and right, respectively). Positive values indicate higher contributions in pCMs, while negative values indicate higher contributions in vCM progenitors. Error bars represent standard deviation (SD). (D-E) Raw sum of contributions for motif syntaxes for *LHX2* (D) and *BMP2* (E). (F) Normalized and Raw sum of contributions for motif syntaxes for *KCNJ3* and *TBX18*.

**Figure S5.**
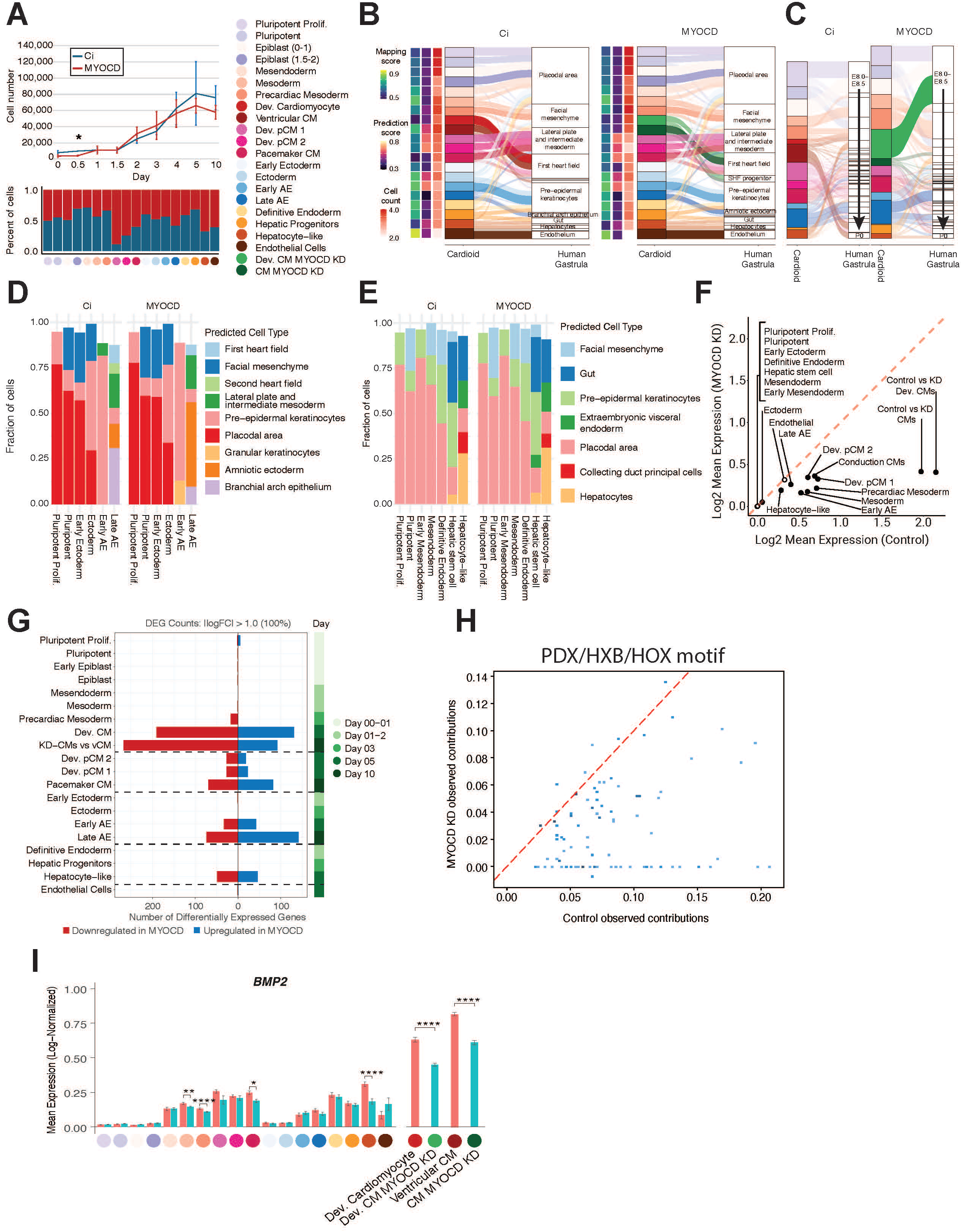
(A) Top - cardioid cell number during differentiation. Astrix marks p-value < 0.05; two-sample two-tailed t-test. Bottom - cell number fraction of Control and *MYOCD* KD cardioids in shared cell-state annotations. (B) The projection breakdown shows the mapping of individual cells to the reference cell states. (C) The projection breakdown shows the mapping of individual cells to the reference developmental stage. (D) Same as Figure 5E but for Late AE. (E) Same as Figure 5E but for Hepatocyte-like cells. (F) *MYOCD* differential expression in the various cell states. (G) Number of DEGs between the control and *MYOCD* KD in all cell states. (H) HOX/PDX/HXB in genome motif contributions in Control versus *MYOCD* KD for all control motif locations. (I) *BMP2* expression across clusters in control and *MYOCD* KD populations. Significance by Wilcox test (* < 0.05; ** < 0.005; **** < 0.0005;).

**Figure S6.**
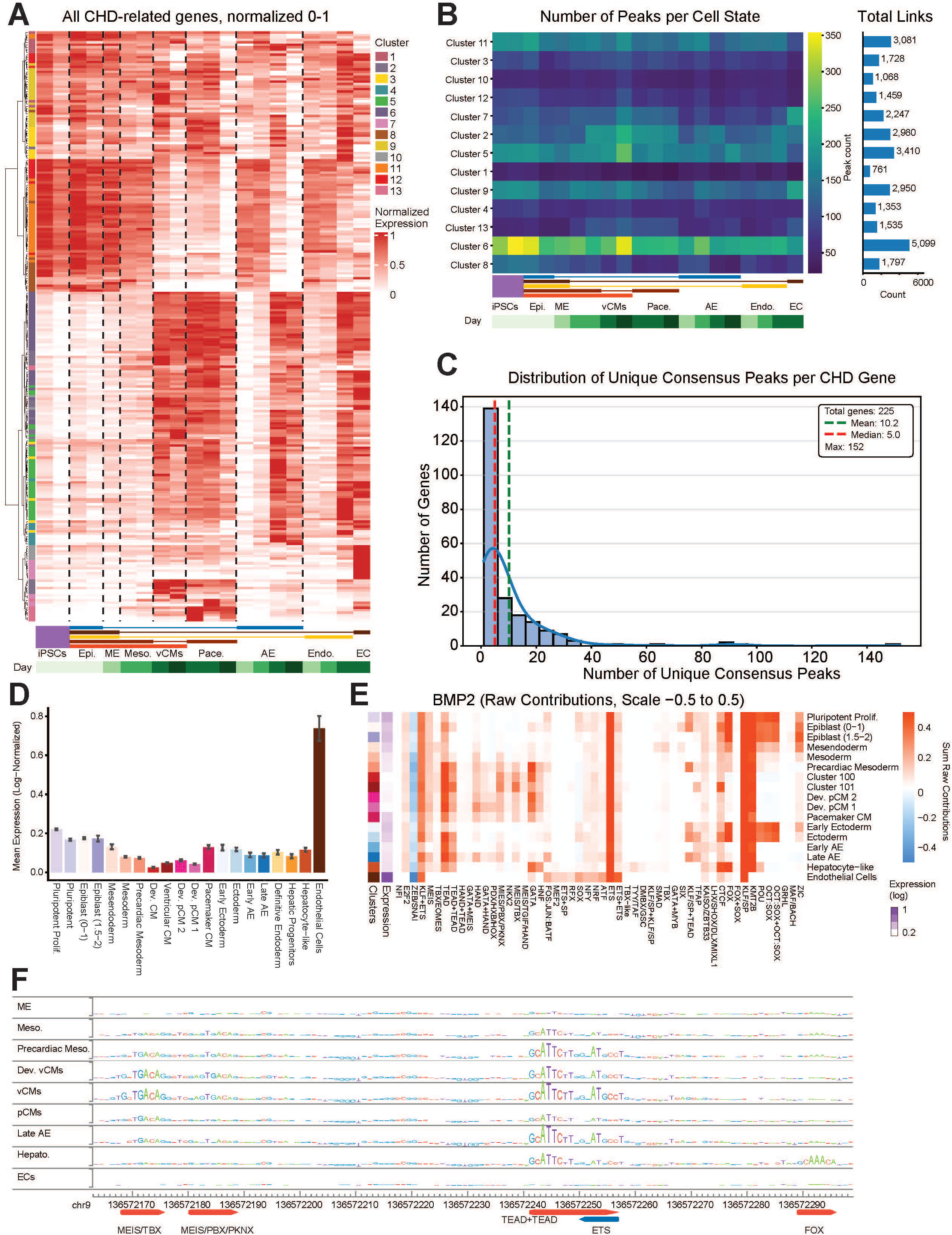
(A) *K*-means cluster breakdown of all CHD genes in Fig. 6A. Hierarchical clustering shows a good correspondence with the k-means clustering (left colored bar). (B) Number of peaks per cell state and the corresponding total number of scE2G links (blue bars) per cell state. (C) Distribution of the number of unique linked consensus peaks per CHD gene across cell states. (D) *NOTCH1* log-norm expression with error bars. (E) Raw sum of contributions for motif syntaxes for *NOTCH1*. (F) cdCRE linked peak to *NOTCH1*, corresponding to Fig. 6D, dashed line.

**Figure S7.**
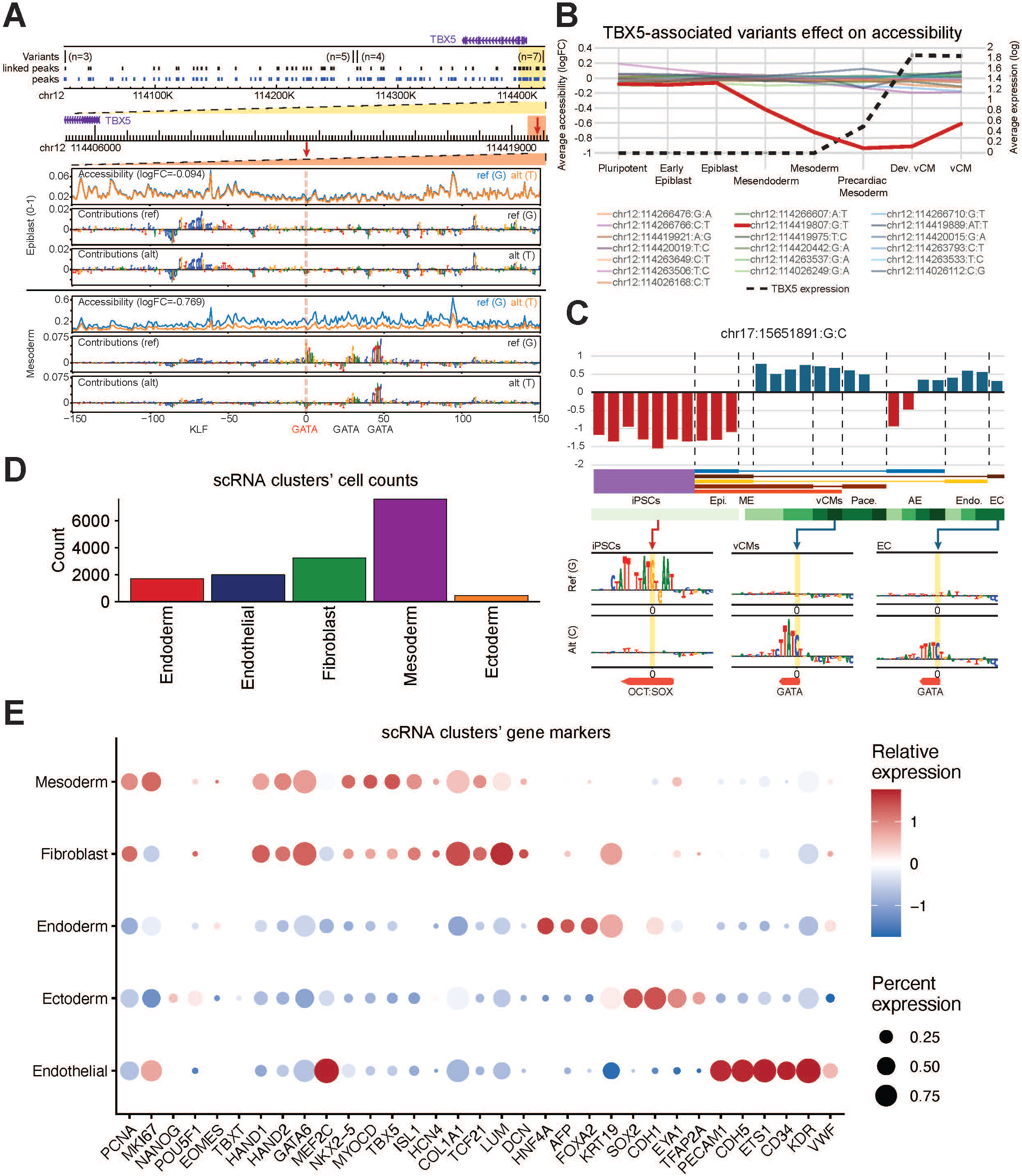
(A) Top - *TBX5* locus and associated variants; Bottom - logFC effect on accessibility and per-base contributions of the linked variant rs377307764 (chr12:114419807:G→C) in Early Epiblast (no effect) and in Mesoderm (logFC = -0.769, AAQ = 0.584). The red dashed line indicates the variant’s position. (B) rs377307764 (chr12:114419807:G→C) accessibility logFC change in the vCM lineage compared to control sequence (left axis). *TBX5* expression (dashed line, right axis). rs377307764 variant in red. (C) Rare variant from gnomAD (MAF 1.46e-5) shows loss of accessibility in early stages of differentiation due to OCT:SOX motifs disruptions. In contrast, the same variant creates a GATA motif in later stages of differentiation across all trajectories. (D) Cell counts in each cluster in both the Control and KI/KI samples. (E) Gene makers for the different clusters.

